# Single-Cell RNAseq Analysis Reveals Robust, Anti-PD-1-Mediated Increase of Immune Infiltrate in Metastatic Castration-Sensitive Prostate Cancer

**DOI:** 10.1101/2022.05.06.490968

**Authors:** Jessica E. Hawley, Aleksandar Z. Obradovic, Matthew C. Dallos, Emerson A. Lim, Karie Runcie, Casey R. Ager, James McKiernan, Christopher B. Anderson, Joel Decastro, Joshua Weintraub, Renu Virk, Israel Lowy, Jianhua Hu, Matthew G. Chaimowitz, Xinzheng V. Guo, Ya Zhang, Jeremy Worley, Mark N. Stein, Andrea Califano, Charles G. Drake

## Abstract

Compared to other malignancies, the tumor microenvironment (TME) of primary and castration-resistant prostate cancer (CRPC) is relatively devoid of immune infiltrates. While androgen deprivation therapy (ADT) induces a complex immune infiltrate in localized prostate cancer, both in animal models and humans, the TME composition of metastatic, castration-sensitive prostate cancer (mCSPC) is relatively unknown and the effects of ADT and other treatments are poorly characterized in this context. To address this challenge, we analyzed metastatic sites from patients enrolled on a phase 2 clinical trial (NCT03951831), in which men were treated with standard-of-care chemo-hormonal therapy with anti-PD-1 immunotherapy, at the single cell level. Longitudinal protein activity-based analysis of TME subpopulations identified immune subpopulations conserved across multiple metastatic sites, their dynamic, treatment-mediated evolution, and associated clinical response features. Our study revealed a therapy-resistant, transcriptionally distinct tumor subpopulation, which comprises an increasing number of cells in treatment-refractory patients, and identified several druggable targets in both tumor and immune cells as candidates to advance treatment and improve outcomes for patients with mCSPC.

## INTRODUCTION

Compared to other tumor types, localized prostate cancer (PC) is characterized by an immunologically ‘cold’ tumor microenvironment (TME), with a relative dearth of immune cells ^1,2^. Preclinical studies in animal models and analyses of human primary PC samples have shown that the small fraction of tumor infiltrating immune cells are tolerogenic and immunosuppressive, as revealed by a dominant fraction of terminally differentiated cytotoxic T cells and regulatory T cells (Treg) ^3–5^. Several studies have shown that androgen deprivation therapy (ADT), the mainstay therapy for advanced prostate cancer, induces immunogenic changes in the TME of castration-sensitive prostate cancer ^6^. This can be attributed to several cooperative mechanisms, including (a) thymic regeneration and increased production of naïve T cells, (b) decreased tolerance and clonal expansion of effector T cells, and (c) stimulation of antigen-specific adaptive immune response. The latter induces robust, chemokines and cytokines secretion-mediated infiltration of functionally competed immune infiltrate into primary prostate tumors ^7–18^. Unfortunately, these pro-immunogenic effects are not durable, and they are further counter-balanced by concomitant increase in immunosuppressive cell subpopulations and decreases in T cell priming ^12,19–21^.

This provides reasonable therapeutic rationale to combine ADT with immunotherapy, thus leveraging anti-tumor immune effects and mitigating ADT-mediated engagement of immune checkpoint activity. To that effect, we designed and initiated the PRIME-CUT Phase II trial (NCT03951831) treating men with metastatic castration-sensitive prostate cancer with a combination of chemo-hormonal therapy and anti-PD-1 immunotherapy, designed to test the combined activity of ADT and a PD-1 inhibitor (cemiplimab) with docetaxel.

This trial helps address the shortcoming that most preclinical and clinical studies have thus far focused on the immunogenic effects of ADT in *primary* prostate cancer. Indeed, the TME of *metastatic*, castration-sensitive prostate tumors and its ADT-mediated remodeling are still elusive and poorly characterized. This is largely due to the challenges associated with longitudinal tissue acquisition from metastatic tumor biopsies, including pre- and post-treatment. In particular, whether the TME of treatment-naïve metastatic sites is “cold”, as for primary prostate cancer, is still largely unknown. Spatial imaging and single-cell sequencing studies have shown a paucity of immune infiltrates in the mCRPC setting, similar to primary PC ^22,23^. By contrast, analysis of PD-L1 protein expression in primary and mCRPC has shown notable differences, with 7.7% and 31.6% of primary PC and mCRPC-derived tissues showing detectable PD-L1 expression, respectively ^24^.

Here, we comprehensively characterized both the tumor and non-tumor subpopulations comprising the mCSPC TME, across distinct metastatic niches—including bone, lymph node, liver, and lung. This was accomplished by leveraging our previously developed pipeline for Virtual Inference of Protein Activity by Enriched Regulons (VIPER) ^25–27^ to study single-cell RNA sequencing (scRNA-seq) profiles of single cells dissociated from these niches. Specifically, akin to a highly-multiplexed gene reporter assay, the differential activity of each protein is assessed based on the differential expression of its lineage-specific transcriptional targets. This allows full activity quantification of ∼ 6,500 proteins in each single cell— including transcriptional regulators and signaling proteins—despite the fact that > 80% of the genes are generally undetected (gene dropout effect). Critically, we have shown that VIPER-based protein activity measurements in single cells compare favorably with antibody-based assays, revealing clinically relevant subpopulations that could not have been detected by gene expression analysis or fluorescence-activated cell sorting (FACS).

Protein activity-based cluster analysis allowed deep stratification of immune and tumor-related subpopulations, virtually eliminating gene dropout for key regulatory and signaling proteins, including critical lineage markers. We applied this methodology to a series of paired, longitudinal metastatic tumor biopsies, including both pre-treatment (baseline) and on-treatment biopsies from eight patients enrolled in the PRIME-CUT Phase 2 clinical trial. In addition to characterizing TME composition differences across patients, we also investigated TME dynamics following ADT therapy—both alone or in combination with anti-PD-1 therapy (prior to docetaxel)—as well as the association between early prostate-specific antigen (PSA) response and the presence of specific TME subpopulations at baseline on a patient-specific basis. Finally, we applied the NY and CA Department of Health approved, CLIA-certified OncoTarget ^28,29^ algorithm for the inference of druggable proteins aberrantly active in single cell subpopulations associated with treatment resistance.

## METHODS

### Study design and participants

The PRIME-CUT study (modulating the <u>Pr</u>ostate cancer Immune <u>M</u>icro<u>e</u>nvironment with <u>C</u>hemoimm<u>u</u>notherapy for metastatic prostate cancer) (Figure 1A), is an open-label, single-arm, phase 2 study (NCT03951831) conducted at Columbia University Irving Medical Center (New York, NY) testing the clinical activity of phased administration of ADT, anti-PD-1 therapy, and docetaxel in men with newly diagnosed metastatic, castration-sensitive prostate cancer. The study was approved by the institutional review board at Columbia University and all participants provided written consent. Key eligibility criteria included diagnosis of metastatic, castration-sensitive prostate cancer with a non-castrate testosterone level (> 150ng/dL). Recruitment was restricted to patients with metastatic lesions amenable to biopsy. Patients with bone metastases were allowed. Treatment consisted of ADT (degarelix 240 mg subcutaneously (SC) for one dose, followed by leuprolide 22.5 mg SC every 12 weeks) followed by anti-PD-1 antibody (cemipliamb-rwlc 350 mg IV every 3 weeks) beginning four weeks after ADT initiation. For potential immune priming, a two-cycle lead-in of anti-PD-1 therapy was administered prior to standard of care docetaxel (75 mg/m^2^ every 3 weeks for six cycles). Participants received ADT and anti-PD-1 antibody until study completion (52 weeks) or until lack of clinical benefit or intolerable side effects. The primary endpoint was the rate of undetectable PSA at 6 months after chemotherapy initiation (37 weeks on-study) compared to the historical rate following ADT plus chemotherapy from the phase III CHAARTED trial ^30^. Secondary endpoints included time to progression to CRPC and rate of radiographic response at study completion. To ensure patient safety within the limitations of a small phase 2 study, toxicity was monitored using a Bayesian method which provides continuous monitoring boundaries for termination of the trial if the toxicity rate is unacceptable ^31^.

**Figure 1:**
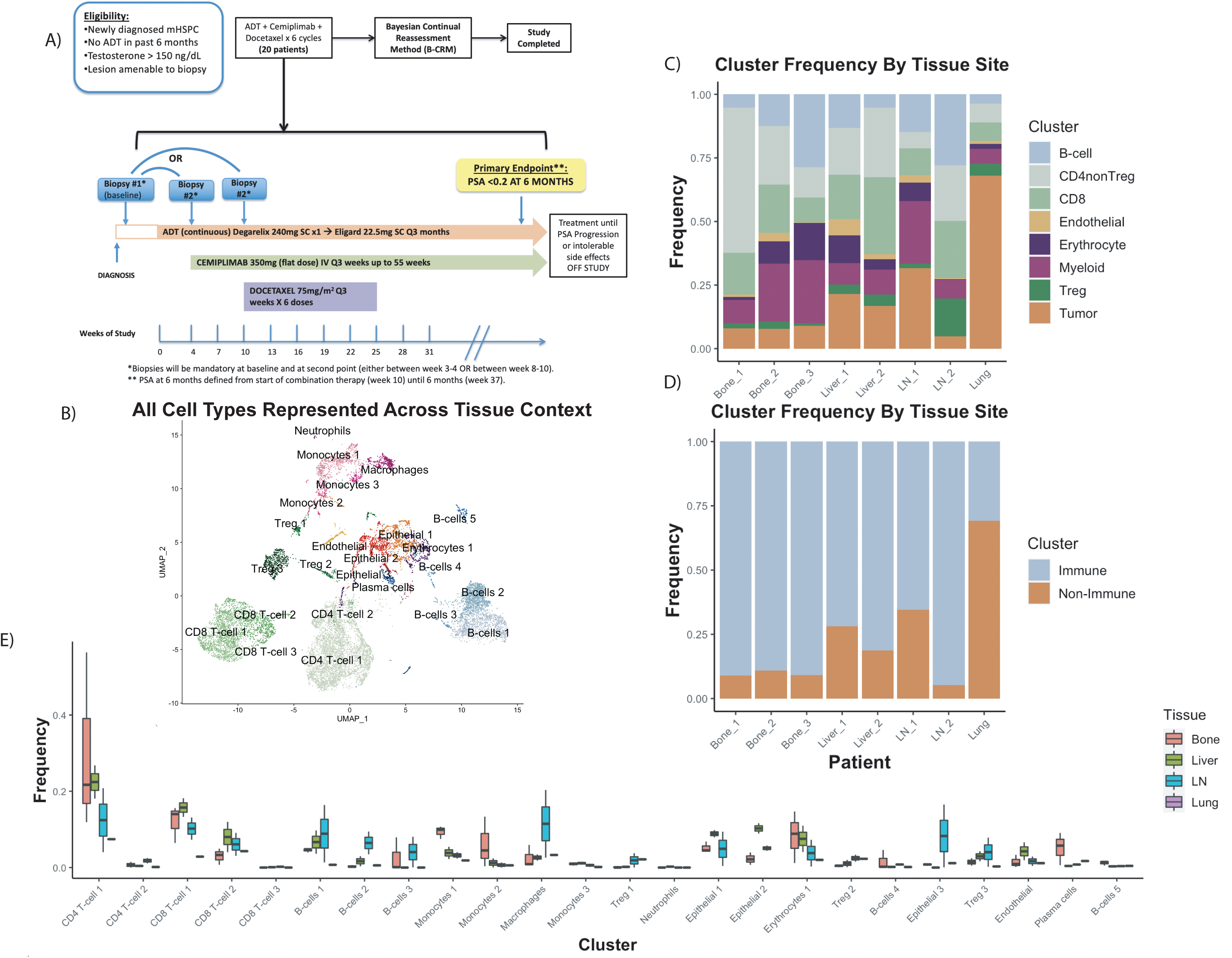
Baseline Composition of Micro-Environment by Tissue Site. A) Phase 2 Trial Design Schema. B) Uniform Manifold Projection (UMAP) plot constructed from VIPER-inferred protein activity of all cells in aggregate across baseline pre-treatment patient samples. Cells are clustered by resolution-optimized Louvain algorithm with cell type inferred by SingleR. C) Stacked barplot of the frequency of each major cell lineage within each baseline patient sample, with each column representing a unique patient and patients grouped by metastatic site. Cell clusters from B are aggregated by shared cell type. D) Stacked barplot of immune vs non-immune cell frequencies, from C. E) Boxplot showing distribution of frequencies for each cell cluster in B at baseline, comparing tissue sites.

### Sample collection

Twelve patients were enrolled on trial from May 2019 through December 2020. At enrollment, all patients were scheduled for an interventional radiology-guided baseline biopsy of the most accessible metastasis. For on-treatment biopsies, subjects were randomized to one of two time points, either 4 weeks or 10 weeks on-study (Figure 1A). Subsequent biopsies at the time of disease progression were optional. All on-treatment biopsies were taken from the same anatomical location as the baseline biopsy. Patients randomized to the week 4 time point were thus treated with four weeks of ADT (degarelix) alone at the time of second biopsy. Patients randomized to the week 10 time point were thus treated with 10 weeks of ADT (4 weeks of degarelix and 6 weeks of leuprolide) as well as two cycles (6 weeks) of anti-PD-1-therapy. We subsequently refer to these time points as ‘ADT only’ and ‘ADT+anti-PD-1,’ respectively. Given the phased administration of ADT and anti-PD-1 therapy, these data allowed us to specifically compare the effects of ADT monotherapy and ADT+anti-PD-1 on the transcriptional program of immune cell subpopulations and prostate tumor cells in the TME across a variety of tissues.

Of the twelve enrolled patients, two patients’ samples were excluded from further analyses due to either an insufficient number of viable cells for loading onto the 10X Genomics instrument or a lack of tumor cells identified in the biopsy sample using copy number inference (see methods below). We thus report on ten patients’ samples from bone, lymph node, liver, and lung metastases. We recovered an adequate number of cells in both baseline and on-treatment biopsy samples from six of the ten patients (four patients with bone metastases, one patient with lymph node metastases, and one patient with lung metastases). In the four remaining patients, only one of the two samples (baseline or on-treatment) per patient yielded adequate cells for sequencing and analysis (one baseline lymph node sample, one baseline liver sample, and one on-treatment bone sample [ADT only], and one on-treatment [ADT+anti-PD-1] liver sample (**Table 1)**.

**Table 1.** Clinical characteristics and cellular yield per biopsy sample.

| Patient | Age | Race | Ethnicity | Baseline<br>PSA<br>(ng/dL) | Biopsy<br>Location | Baseline<br>(no.<br>cells) | ADT only<br>(no. cells) | ADT +<br>anti-PD-1<br>(no. cells) | Recurrence<br>(no. cells) |
| --- | --- | --- | --- | --- | --- | --- | --- | --- | --- |
| 1 | 62 | White | Not<br>Hispanic or<br>Latino | 11.56 | Bone | 2360 | NA | 686 | NA |
| 3 | 65 | White | Not<br>Hispanic or<br>Latino | 156.4 | Bone | 1027 | 995 | NA | 764 |
| 5 | 82 | Black | Other<br>Hispanic or<br>Latino | 26.2 | LN | 4090 | 5153 | NA | NA |
| 6 | 52 | White | Dominican | 11.93 | Liver | 0 | NA | 2475 | NA |
| 7 | 60 | Other | Dominican | 146.7 | Lung | 699 | NA | 1956 | NA |
| 8 | 49 | White | Not<br>Hispanic or<br>Latino | 91.93 | Bone | 606 | 1784 | NA | NA |
| 10 | 76 | White | Dominican | >5000 | Bone | 521 | NA | 97 | NA |
| 12 | 65 | Black | Dominican | 95.05 | Bone | 0 | 913 | NA | NA |
| 13 | 54 | Other | Dominican | 229.9 | LN | 1212 | NA | 0 | NA |
| 14 | 63 | Black | Not<br>indicated | 90.71 | Liver | 2161 | 0 | NA | NA |

LN, lymph node; no., number; NA, not applicable; PSA, prostate specific antigen.

### Tissue Dissociation

Fresh tumor was minced to 2-4 mm sized pieces with micro-scissors. For bone metastases, minced tissue was resuspended and examined under microscope. If already found to be dissociated to a single-cell suspension, the entire sample was passed through a 70-um filter for downstream processing. For bone metastases not found to be dissociated and for all other tissue sites, tissue was digested with 2.5 mL of the tumor dissociation medium (L-15 medium with 1g/L glucose, 5% FBS, 15 mM HEPES, 800 U/ml collagenase IV, and 0.1 mg/ml DNase I) with gentle agitation in a glass vial in 37°C water bath for 30 min. Dissociated cells were passed through a 70-um filter and centrifuged at 300 g for 5 min at 4°C. If the pellet was red or pink, the cells were incubated with ACK lysis buffer and centrifuged again at 300 g for 5 min at 4°C. Cells were resuspended in L-15 medium with 1g/L glucose, 5% FBS, and 15 mM HEPES and counted with Trypan Blue for viability assessment. An aliquot of 10,000-20,000 cells per sample were used for subsequent single-cell RNA sequencing analysis.

### Single-cell RNA sequencing and data processing

Single-cell RNA sequencing (scRNA-seq) analysis was performed using the 10x Genomics (Pleasanton, CA) Chromium Single Cell 5’ Library & Gel Bead Kit at the Columbia University Human Immune Monitoring Core (HIMC). Manufacturers’ protocols were followed for the preparation of gene expression libraries and the subsequent sequencing on the Illumina (San Diego, CA) NovaSeq 6000 Sequencing System. The sequencing reads (base call files) were converted to FASTQ files using the 10x Genomics data processing pipeline “cellranger mkfastq”, followed by the “cellranger count” for cell calling, gene mapping to the pre-built human reference set of 30,727 genes (10x Genomics), and gene counting. The cell-gene count matrix data were then processed using the publicly available Seurat package to filter for cells with less than 10% mitochondrial RNA content, more than 1,500 UMI counts, and fewer than 15,000 UMI counts, followed by the Seurat SCTransform command to perform a regularized negative binomial regression based on the 3000 most variable genes. Samples were then combined by the Seurat Anchor Integration algorithm. To ensure statistically valid clustering, the resulting matrix was clustered using the Louvain Algorithm, with resolution selected automatically to maximize clustering silhouette score, as previously described ^27^. Gene Expression data were projected into their first 50 principal components using the RunPCA function in Seurat, and further reduced into a 2-dimensional visualization space using the RunUMAP function with method umap-learn and Pearson correlation as the distance metric between cells. Differential Gene Expression between clusters was computed by the MAST hurdle model for single-cell gene expression modeling, as implemented in the Seurat FindAllMarkers command, with log fold change threshold of 0.5 and minimum fractional expression threshold of 0.25, indicating that the resulting gene markers for each cluster are restricted to those with log fold change greater than 0 and non-zero expression in at least 25% of the cells in the cluster.

### Semi-Supervised Cell Type Calling

For each single cell gene expression sample, cell-by-cell identification of cell types was performed using the SingleR package ^32^ and the preloaded Blueprint-ENCODE reference, which includes normalized expression values for 259 bulk RNASeq samples generated by Blueprint and ENCODE from 43 distinct cell types representing pure populations of stroma and immune cells ^33,34^. The SingleR algorithm computes correlation between each individual cell and each of the 259 reference samples, and then assigns both a label of the cell type with highest average correlation to the individual cell and a p-value computed by Wilcox test of correlation to that cell type compared to all other cell types. Cell-by-cell SingleR labels were restricted to those with *p* <0.05, and unsupervised clusters are labelled as a particular cell type based on the most-represented SingleR cell type label within that cluster. Since tumor cells are not represented within the Blueprint-ENCODE reference, tumor cells are typically assigned as ‘epithelial,’ since prostate cancer is epithelial in origin. The tumor cell identity of these cells was further confirmed by expression of KLK3, a prostate cancer marker gene, as well as by inference of copy number variations using the InferCNV algorithm ^35^ with all lymphoid and myeloid cell clusters specified as a copy-number-normal reference.

### Regulatory network and protein activity inference

Protein activity was measured from single-cell gene expression profiles according to the pipeline previously described ^36^ and subsequently used for analysis of single-cell ccRCC samples ^27^. From the combined and batch-corrected dataset of all patients, metaCells were independently assembled from each distinct single cell subpopulation, as identified by gene expression clustering. Specifically, metaCells were generated by summing SCTransform-corrected template counts for the 10 nearest neighbors of randomly selected cells, based on Pearson correlation distance in gene expression space. 200 cells were randomly sampled from each cluster to generate metaCells and the latter were then used to generate cluster-specific regulatory networks using the ARACNe algorithm. ARACNe was run with 100 bootstrap iterations using 1,785 transcription factors (genes annotated in gene ontology molecular function database as GO:0003700, “transcription factor activity”, or as GO:0003677, “DNA binding” and GO:0030528, “transcription regulator activity”, or as GO:0003677 and GO:0045449, “regulation of transcription”), 668 transcriptional cofactors (a manually curated list, not overlapping with the transcription factor list, built upon genes annotated as GO:0003712, “transcription cofactor activity”, or GO:0030528 or GO:0045449), 3,455 signaling genes (annotated in GO biological process database as GO:0007165, “signal transduction” and in GO cellular component database as GO:0005622, “intracellular” or GO:0005886, “plasma membrane”), and 3,620 surface markers (annotated as GO:0005886 or as GO:0009986, “cell surface”). ARACNe was used to analyze each of these gene sets independently thus ensuring that the data processing inequality (DPI) was not affected by different baseline mutual information for different protein classes. Moreover, we did not use ARACNe to infer the regulatory targets of proteins with no known signaling or transcriptional activity, for which the effect on downstream transcriptional targets may be hard to interpret. Parameters were set to zero DPI tolerance and Mutual Information (MI) p-value threshold of *p* = 10^-8^, computed by shuffling the original dataset as a null model. Protein activity was inferred by VIPER using the cluster-specific ARACNe networks on the SCTransform-scaled and Anchor-Integrated gene expression signature of all single cells from each patient. The resulting protein activity matrix was loaded into a Seurat Object with CreateSeuratObject, then projected into its first 50 principal components using the RunPCA function, and further reduced into a 2-dimensional visualization space using RunUMAP function with method umap-learn and Pearson correlation as the distance metric between cells. Differential protein activity between clusters identified by resolution-optimized Louvain was computed using Student’s t-test, and top proteins for each cluster were ranked by p-value.

### Association of TME subpopulations with tissue site and PSA response

Cell counts per cluster were normalized within each individual sample to cluster frequencies, and subsequent comparisons were made between cluster frequencies in different tissue sites, treatment time-points, and at baseline between patients who responded or did not respond to treatment, as assessed by change in PSA over time (Figure 4). This was done for clusters identified by the Louvain algorithm in combined dataset of all cells, representing the entire TME. Separately, tumor cell clusters were isolated as a new Seurat object on which principal components and UMAP projection were re-computed from the VIPER-inferred protein activity matrix. These were subsequently sub-clustered by resolution-optimized Louvain algorithm ^27^. Differential protein activity was computed for each tumor cell subcluster, by Student’s t-test, with results shown in Figure 6, and pathway enrichment within each cluster was assessed by the Enrichr browser tool ^37^ (Figure S5). Tumor cell counts within each subcluster were normalized to the total count of all tumor cells to compare relative frequencies of each tumor cell population at baseline in patients with early response to treatment (defined by reduction to less than 1% of initial PSA within 10 weeks of treatment) compared to patients with initial response followed by PSA progression (late progressors). The same was done to compare frequencies of each population by tissue site and by treatment time-point.

### Tumor cell subcluster OncoTarget analysis and association with outcome in external datasets

Druggable protein activity within tumor cell subclusters was evaluated by the OncoTarget algorithm ^28,29^, in which the log(UMI/million) gene expression of each tumor cell was scaled by z-score against the log(TPM) gene expression of the entire TCGA database as a reference, and VIPER was applied using cluster-specific ARACNe networks. From the resulting protein differential activity matrix, a subset of proteins was selected for which a high-affinity inhibitor already exists, as determined by analysis of the DrugBank online database ^28^. The activity scores of each druggable protein, as computed on a cell-by-cell basis in each distinct cluster, were transformed into p-values by fitting to the analytical normal distribution; for each cell multiple hypothesis testing correction was implemented using Bonferonni’s method. Corrected p-values were converted to S = -log10(p-value) (scores) for ease of visualization, and only proteins with mean S ≥ 5 in each cluster were reported. The resulting druggable protein activity matrix is shown in Figure 7.

For each transformed subpopulation, a protein signature was defined based on the proteins that were significantly differentially activated in that cluster. Then for each of three independent external prostate cancer bulk-RNA-Seq datasets, (TCGA, East Coast SU2C, West Coast SU2C) ^38–40^, enrichment of these protein signatures was assessed as follows. First, the bulk-RNA-Seq dataset was internally scaled by z-score, then VIPER protein activity inference was performed using the single-cell ARACNe networks, and finally enrichment of each tumor subcluster signature was determined in each bulk-RNA-Seq sample by Gene Set Enrichment Analysis (GSEA) ^41^, where genes were ranked from the highest to lowest activity of their encoded protein. The resulting normalized enrichment scores were tested against recurrence-free-survival time in TCGA or overall survival time in SU2C by Cox regression (Figure 8A). Since enrichment of tumor cell cluster 1 was found to be significantly associated with shorter recurrence-free-survival in TCGA, the genes in the leading-edge, and their encoded proteins, were further identified by GSEA analysis of all proteins ranked by differential activity in TCGA samples with and without recurrence. Activity of these proteins for all TCGA samples is shown in Figure 8B. Finally, patient-by-patient enrichment scores were binarized to less than zero = “low” and greater than zero = “high” and assessed for effect on survival by log-rank test and Kaplan-Meier curve, with all statistically significant results shown in Figure 8C-F.

### Quantification and Statistical Analysis

All quantitative and statistical analyses were performed using the R programming environment and packages described above. Differential gene expression was assessed at the single-cell level by the MAST single-cell statistical framework as implemented in Seurat v3, and differential VIPER activity was assessed by t-test, each with Bonferroni multiple-testing correction. Comparisons of cell frequencies were performed by non-parametric Wilcox rank-sum test, and survival analyses were performed by log-rank test and cox regression. In all cases, statistical significance was assessed at *p* ≤ 0.05. Details of all statistical tests used can be found in the corresponding figure legends.

## RESULTS

### Gene expression and protein activity clustering reveals a robust immune infiltrate in metastatic castration-sensitive prostate cancer

Since primary prostate cancer (PC) is characterized by a relative immune desert, with low representation of tumor-infiltrating immune cell subpopulations ^1,2^, we tested whether the tumor microenvironment (TME) of metastatic, castration-sensitive prostate cancer (mCSPC) patients was similarly immunologically ‘cold’. For this purpose, we collected pre-treatment (baseline) metastatic needle-core biopsies from 8 patients (Table 1), across 4 different metastatic niches (bone, lymph nodes, liver, and lung). We then dissociated live cells to generate scRNA-seq profiles, which were then first used to perform gene expression-based cluster analysis. Cell lineages were inferred for each single-cell using the SingleR algorithm ^32^, with clustering performed using resolution-optimized Louvain ^27^. Standard gene expression-based clustering revealed 15 overall clusters across all metastatic sites, corresponding to 12 distinct immune cell subpopulations, as well as subpopulations comprising fibroblasts, endothelial, and epithelial cells (Figure S1). Inspection of the top five most differentially upregulated genes in each cluster (Figure S2) was consistent with the ascribed cellular identities assigned by SingleR. For example, granzyme M (GZMM) and natural killer granule 7 (NKG7) were differentially upregulated in CD8 T cells, and CD37 in B cells. SingleR does not provide classification of tumor vs. normal cells; as a result, tumor cells with an epithelial origin, such as prostate cancer cells, were labelled as ‘epithelial cells.’ The tumor-cell identity was then confirmed based on the expression of tumor marker genes, such as KLK3, and by inferred Copy Number Variation analysis (Figure S6).

Due to high gene dropout levels, scRNA-seq analyses can typically monitor ∼20% of all genes in each cell ^27^. To address this issue, we relied on an established single cell analysis pipeline, which leverages the VIPER algorithm to measure protein activity from single-cell gene expression data. By assessing protein activity based on the expression of 100 transcriptional targets of each protein, VIPER dramatically mitigates gene dropout effects, thus allowing even detection of proteins whose encoding gene was completely undetected ^27^. Protein activity-based clustering revealed a much finer-grain subpopulation structure, comprising 24 distinct subpopulations, primarily comprised of immune cells but also including erythrocytes, endothelial, and three molecularly distinct transformed epithelial cell clusters (Figure 1B). In particular, while gene expression-based cluster analysis yielded a single homogeneous subpopulation of monocytes and macrophages, activity-based analysis stratified these cells into five subpopulations, representing three distinct monocyte subtypes, as well as macrophages and neutrophils. There was also further refinement of T cells, resulting in the identification of subpopulations comprising T-regulatory (Treg) and CD8 T cells. Furthermore, subpopulations comprising five B cell and one plasma cell subtypes were identified by VIPER analysis, compared to only one and two subtypes, respectively, by gene expression. Critically, VIPER analysis identified three distinct subpopulations of transformed epithelial cells, while only a single cluster could be identified by gene expression analysis. This is critical as these subpopulations may represent distinct drug sensitivity mechanisms.

Overall, the mean proportion of immune cells across all metastatic sites was 87% (range: 30.9% [lung] – 94.6% [lymph node]) thus far exceeding the sparse immune infiltration typically seen in primary prostate cancer ^5,42^. While this degree of immune infiltration may seem surprising, the data are broadly consistent with prior warm-autopsy findings in advanced castrate-resistant prostate cancer ^43^, where CD14+/CD206+ macrophage proportions ranged from 10-50% in metastatic lesions of the lymph node, dura mater, liver, bone marrow, and adrenal glands. Prior work did not assess T-cell infiltration, but the reported levels of myeloid infiltration are in line with our observations here. While cryopreservation in tissue processing has been shown to deplete epithelial cells, leading to potential methodological bias, no such bias has been previously noted from freshly dissociated tissue, as collected in this study ^44^. However, since the primary goal of this study was to assess relative differences in tumor micro-environment composition across tissue contexts and in response to treatment, any artefactual depletion of non-immune cells is expected to be consistent across samples, thus preserving the validity of subsequent inter-sample comparisons.

### Protein activity analyses show distinct differences in immune cell subpopulations across metastatic sites

The proportional representation of immune cell subtypes in the TME varies broadly, depending on tissue type ^45,46^. To compare subtype composition of the TME across different prostate cancer metastatic sites, we collapsed the initial 24 VIPER clusters into eight lineage-specific meta-clusters, including B cells, CD4 non-Treg, CD8, endothelial, erythrocyte, myeloid, Treg, and tumor cells. We then visualized the relative representation of these coarse-grain subpopulations across the four metastatic niches, prior to ADT or anti-PD-1 treatment (Figure 1C, 1D). The baseline biopsy from the lung sample was comprised mostly of tumor cells, with associated CD4, CD8 and myeloid populations representing only 30.9% of all cells (Figure 1C). By contrast, the liver biopsies samples contained a much higher proportion of immune cells, representing 77.9% of all cells, on average. Since lymph nodes are part of the primary immune system, it was perhaps unsurprising to find that, on average, immune cells comprised 94.6% of all cells in these samples. Finally, also consistent with their niche composition, bone lesions were also highly enriched in immune cells, which represented 90.5% of all cells on average.

We next compared the frequency of the 24 different cellular subpopulations identified across the four distinct metastatic sites (*i.e.*, bone, lymph node, liver, and lung) (Figure 1E), whose most significantly differentially activated proteins (subpopulation markers) are shown in Figure 2. In bone-derived metastases, as expected, plasma cells were highly enriched relative to other sites (*p* ≤ 0.05). Additionally, there was an increased representation of monocyte 1 and 2 subtypes, relative to other metastatic sites, with increased activity of transcriptional repressors (*e.g.*, BATF3), transcriptional activators (*e.g.*, SH3BP2), regulators of G-protein signaling (*e.g.*, RGS18), and serine proteases (*e.g.*, PRTN3). We also noted significant over-representation of erythrocytes in bone metastases (*p* ≤ 0.05) with high activity of epithelial cell transforming 2 (ECT2), Rho GTPase Activating Protein 11A (ARHGAP11A), and Kinesis Family Member 14 (KIF14), which play established roles in mitosis, cell-cycle arrest, and microtubule motor proteins respectively ^47–49^. These likely represent a population of dividing erythroid progenitor cells captured incidentally during bone marrow biopsy. In lymph node samples, a robust B cell population was detected (B cell 2). There was also an increased proportion of T regulatory cells (Treg 3), with elevated activity of TNFSRF18 (GITR), in the lymph nodes compared to other metastatic sites (*p* ≤ 0.05). Of interest, this specific T regulatory population had high activity levels of ETS Variant Transcription Factor 1 (ETV1) (*p* ≤ 0.05), a gene known to be overexpressed in prostate cancer ^50,51^. To our knowledge, this has not been previously described in T regulatory cells of prostate cancer tumor metastases and supports recent findings that immune cells may mimic expression of tumor marker genes ^52^. Liver metastases had immune infiltrations similar to bone metastases, in both overall proportion and subpopulation frequency. Notably across all tissues there was a large proportion of CD8 T cells (CD8 T cell 1 and 2) and CD4 non-Treg T cells (CD4 T cell 1). The CD8 T cell 2 cluster was chiefly defined by increased protein activity of lymphocyte activation gene 3 protein (LAG3), an inhibitory immune receptor ^53^. Finally, the single lung metastasis profiled was notably the least immune-infiltrated at baseline, with only 30.9% immune cells overall.

**Figure 2:**
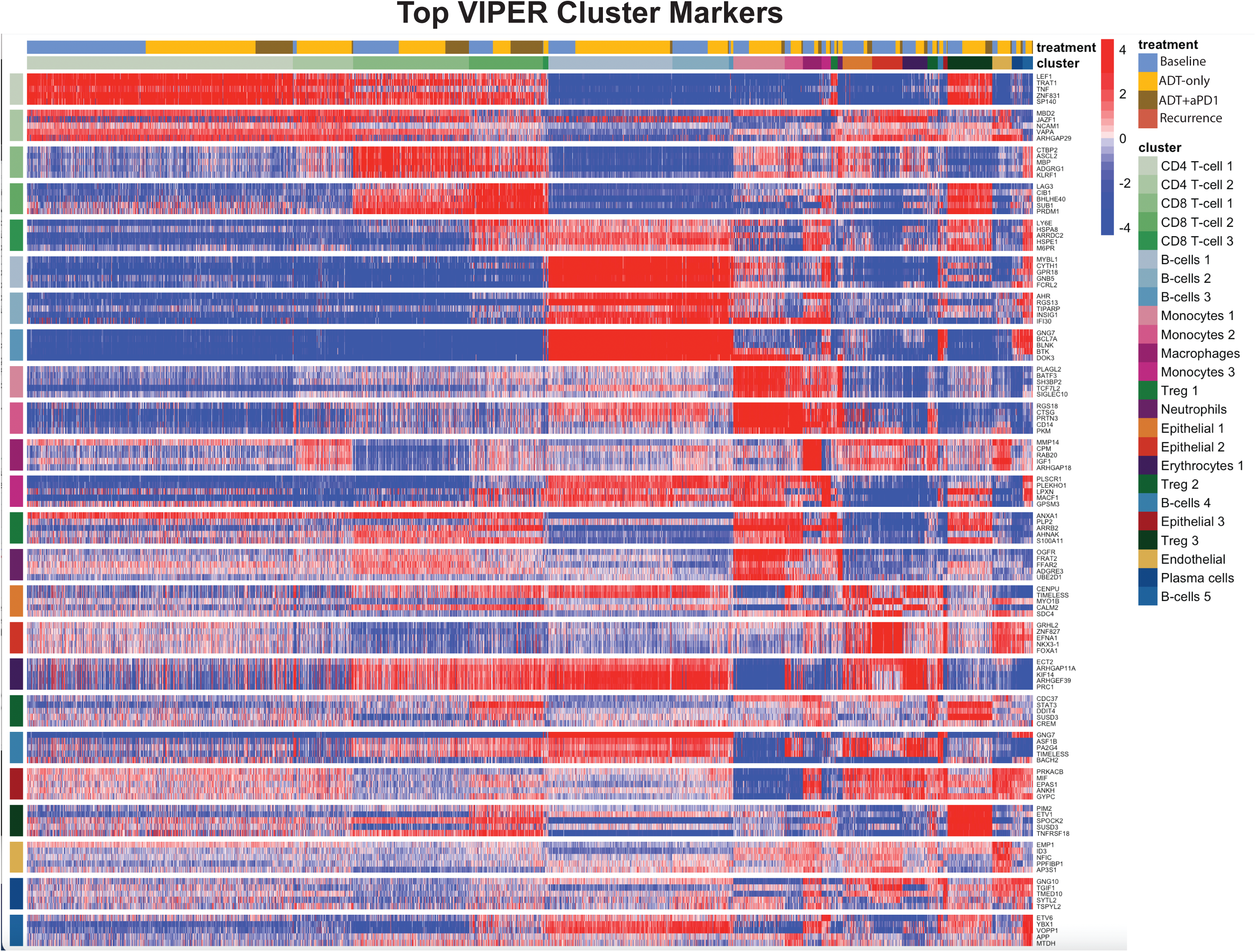
Top Protein Activity Cell Cluster Markers. Heatmap of top 5 most differentially activated proteins for each cell type cluster from aggregate single-cell RNA-Sequencing data across all patient samples. Each row represents a protein, grouped by cluster in which they are the most active, with cluster labels on the x and y-axes. Each column represents a single cell. Above the x-axis cluster label there is also a treatment label indicating timepoint at which a given cell was profiled.

### Treatment with combination ADT plus anti-PD-1 results in a significant expansion of CD8 T cells across several metastatic sites

To compare the immunologic effects of ADT on the TME dynamics of the four different metastatic sites, either alone or in combination with the anti-PD-1 cemiplimab, we compared the pre- vs. post-treatment frequency of each subpopulation across the four metastatic sites, see UMAP plots and stacked bar graphs (Figure 3). As discussed, all patients on trial were required to have a baseline metastatic biopsy, as well as an on-treatment biopsy, with patients randomized to one of two time points for the on-treatment biopsy, at either four weeks after ADT (degarelix) initiation or after ADT plus two cycles of cemiplimab (anti-PD1). Overall, sufficient patients with bone and lymph node metastases were enrolled to enable collection of biopsy samples at baseline and at both on-treatment time points. Additionally, we were also able to obtain a tumor progression biopsy from a patient with subsequent tumor recurrence in the bone, after 11 months on treatment. All liver and lung metastatic biopsy samples were collected at baseline and after ADT with two cycles of anti-PD-1. No samples of liver and lung metastases were collected after ADT alone given the randomization procedures based on patients’ order of enrollment.

**Figure 3:**
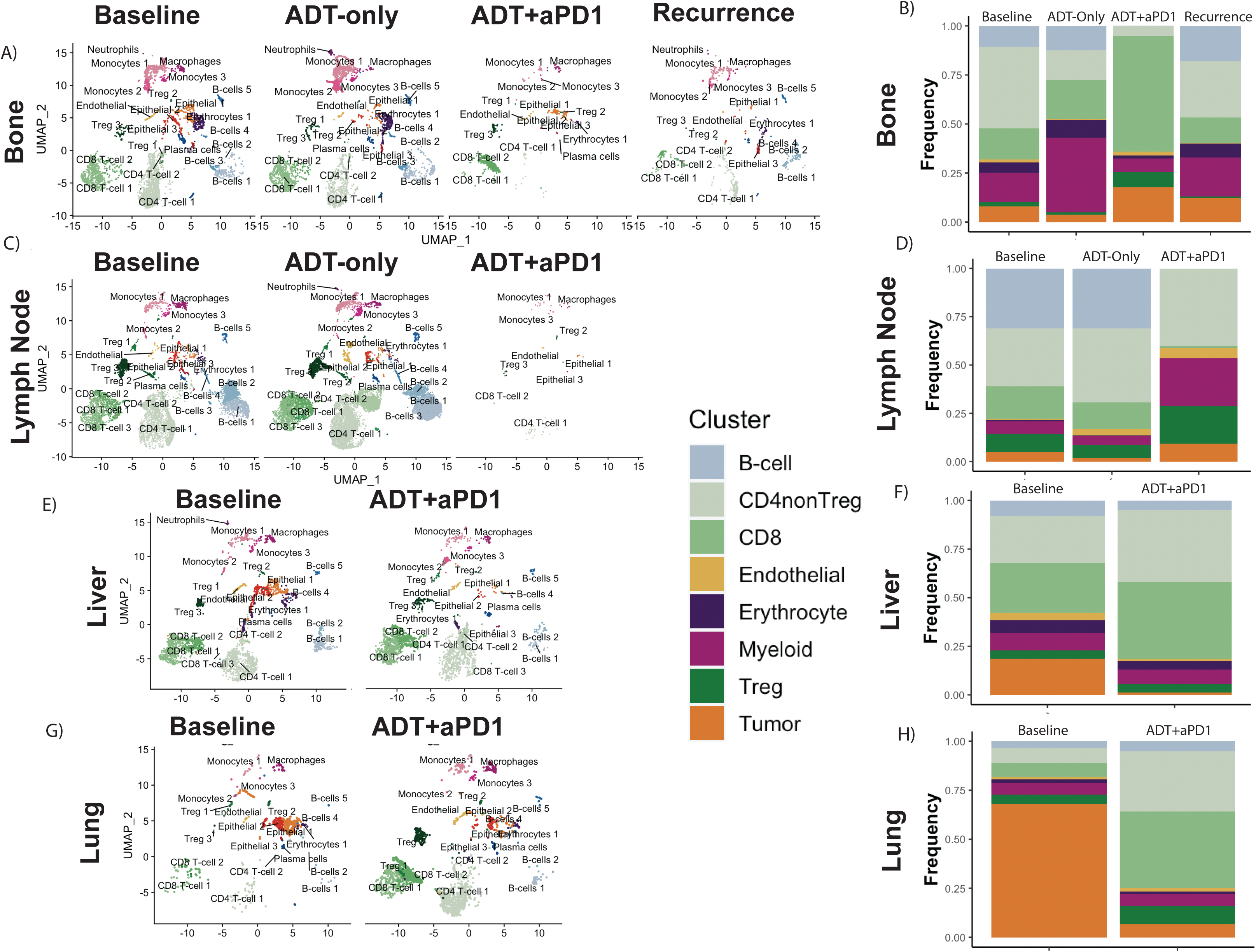
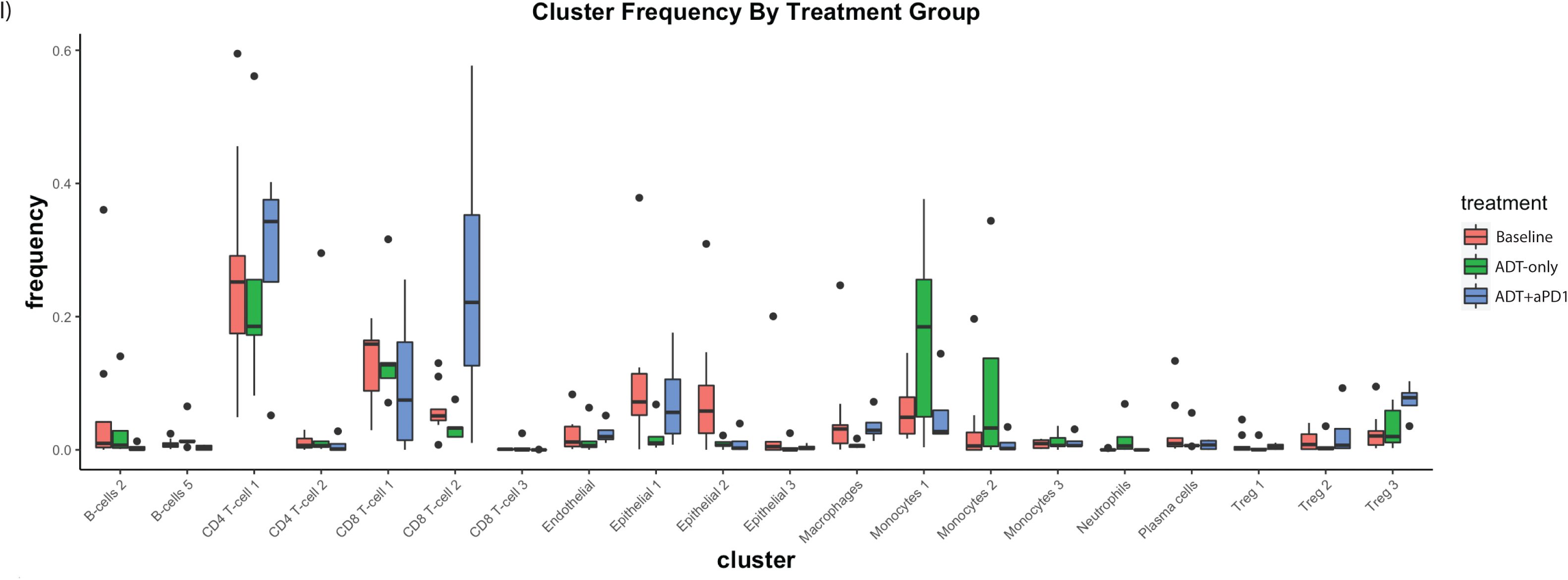
Treatment with ADT+anti-PD-1 Induces Dramatic Changes in the Tumor Micro-Environment. A) UMAP plot of all cells from patients with metastatic Bone lesions, split by treatment time-point (Baseline, ADT-only, ADT+anti-PD-1, and post-treatment Recurrence) and labelled by cell cluster. B) Stacked barplot showing the relative frequency of each major cell lineage by treatment time-point for patients with metastatic Bone lesions, with each column representing aggregate of all samples profiled at a specific treatment time-point. C) UMAP plot, as in A, for patients with metastatic Lymph Node lesions. D) Stacked barplot, as in B, for patients with metastatic Lymph Node lesions. E) UMAP plot, as in A, for patients with metastatic Liver lesions. F) Stacked barplot, as in B, for patients with metastatic Liver lesions. G) UMAP plot, as in A, for patients with metastatic Lung lesions. H) Stacked barplot, as in B, for patients with metastatic Lung Lesions. I) Boxplot showing distribution of frequencies for each cell cluster, comparing frequencies across treatment time-points including Baseline, ADT-only, and ADT+anti-PD-1.

Treatment-mediated evolutionary pressure can induce complex changes in tumor cells and in the TME. We thus leveraged pre- and on-treatment scRNA-seq profiles to interrogate the ADT and ADT plus anti-PD-1-mediated TME dynamics. Considering the coarser subpopulation representation—including B-cells, CD4 non-Tregs, Tregs, CD8 T-cells, Myeloid cells, Endothelial cells, Erythrocytes, and Tumor cells—we characterized treatment-mediated changes at each metastatic site. In bone-derived samples, consistent with pre-clinical data ^12^, ADT induced increased representation of myeloid cells (*p* = 2.8e-118 by Fisher’s Exact Test), while decreasing CD4 non-Treg cells and tumor cells (*p* = 0.01 and 1.1e-15) (Figure 3A-B). By contrast, the ADT and anti-PD-1 combination reduced myeloid cell representation (*p* = 3.6e-10) while concomitantly increasing CD8 T cells (*p* = 3.1e-115). In the single progression biopsy from a bone lesion, the relative subpopulation frequencies resembled that of the baseline samples, albeit with a greater proportion of tumor cells. Finally, in lymph node-derived samples, ADT induced mild expansion of CD4 non-Treg cells (*p* = 5.3e-36) (Figure 3C-D) even though, contrary to bone-derived samples, myeloid subpopulations were not expanded. These findings are consistent with the notion that treatment-induced immunologic changes vary based on the metastatic niche.

Few cells were recovered from lymph nodes after combination treatment with of ADT and anti-PD-1. However, the recovered cells comprised a greater proportion of Treg cells (*p* = 0.002) and myeloid cells (*p* = 1.4e-8), with virtually no representation of CD8 T-cells or B cells. Surprisingly, in both the bone and lymph node-derived samples, there was a relative increase in tumor cell composition following combination therapy with ADT and anti-PD-1, compared to baseline and ADT-only samples. This contrasts with observations from the viscera (liver and lungs), where combination therapy demonstrated substantial reduction in overall tumor cell composition (*p* = 4.1e-119 in liver, *p* = 4.0e-222 in lungs) (Figure 3E-H). Additionally, the myeloid compartment expansion observed in bone-derived samples (after ADT) and in lymph node-derived samples (after combination therapy) was not detected in the viscera (liver and lungs). In contrast, similar to bone-derived samples, combination therapy induced significant increase in CD8 T cells in liver and lung-derived samples (*p* = 1.6e-29 and 7.5e-66, respectively), which was not observed with ADT alone. Taken together, these data show that anti-PD-1 immunotherapy increased CD8 T cell infiltration into metastatic sites, when used in combination with ADT, to an extent that was not observed with ADT alone.

We next focused on the finer-grain subtypes detected by protein activity analysis. Specifically, we assessed which subtypes showed treatment-mediated relative composition increase or decrease (Figure 3I). After ADT treatment, the fraction of Cluster 2 CD8 T cells decreased relative to baseline (*p* = 0.034 by Mann-Whitney U-test), while all monocyte subtypes increased, across all metastatic sites (*p* = 0.036). In contrast, following combination therapy, we observed significant expansion of Cluster 1 CD4 T cells (characterized by high TNF activity; Figure 2), Cluster 2 CD8 T cells (characterized by high LAG3 activity ; Figure 2), and Cluster 3 Treg cells (characterized by high TNFRSF18 activity; Figure 2), across all metastatic sites (*p* = 0.033, 0.026, and 0.008, respectively). These three populations represent the bulk of tumor-infiltrating immune cells whose composition is increased by anti-PD-1 therapy and are consistent with the expected effects of PD-1 blockade on the T cell compartment ^54^.

### Baseline Immune Cell Population Frequencies Associate with Treatment Response

We next sought to associate baseline subpopulation composition with treatment response, with the goal of determining whether the presence of specific pre-treatment subpopulations may be associated with PSA response or progression after therapy. To that end, we categorized patients into three treatment response groups (early PSA response, stable disease, or late progressors) based on PSA log10 fold-change (Figure 4A). Thus, early responders showed PSA decrease below 1% of the pre-treatment value, indicating robust response, while late progressors initially responded to therapy, with marked PSA decrease, but showed PSA increase by week 28. We then compared the frequencies of each of the 24 lineage subtypes in ‘early responders’ vs. ‘late progressors’. To mitigate confounding factors, patients with stable disease were excluded from the analysis.

**Figure 4:**
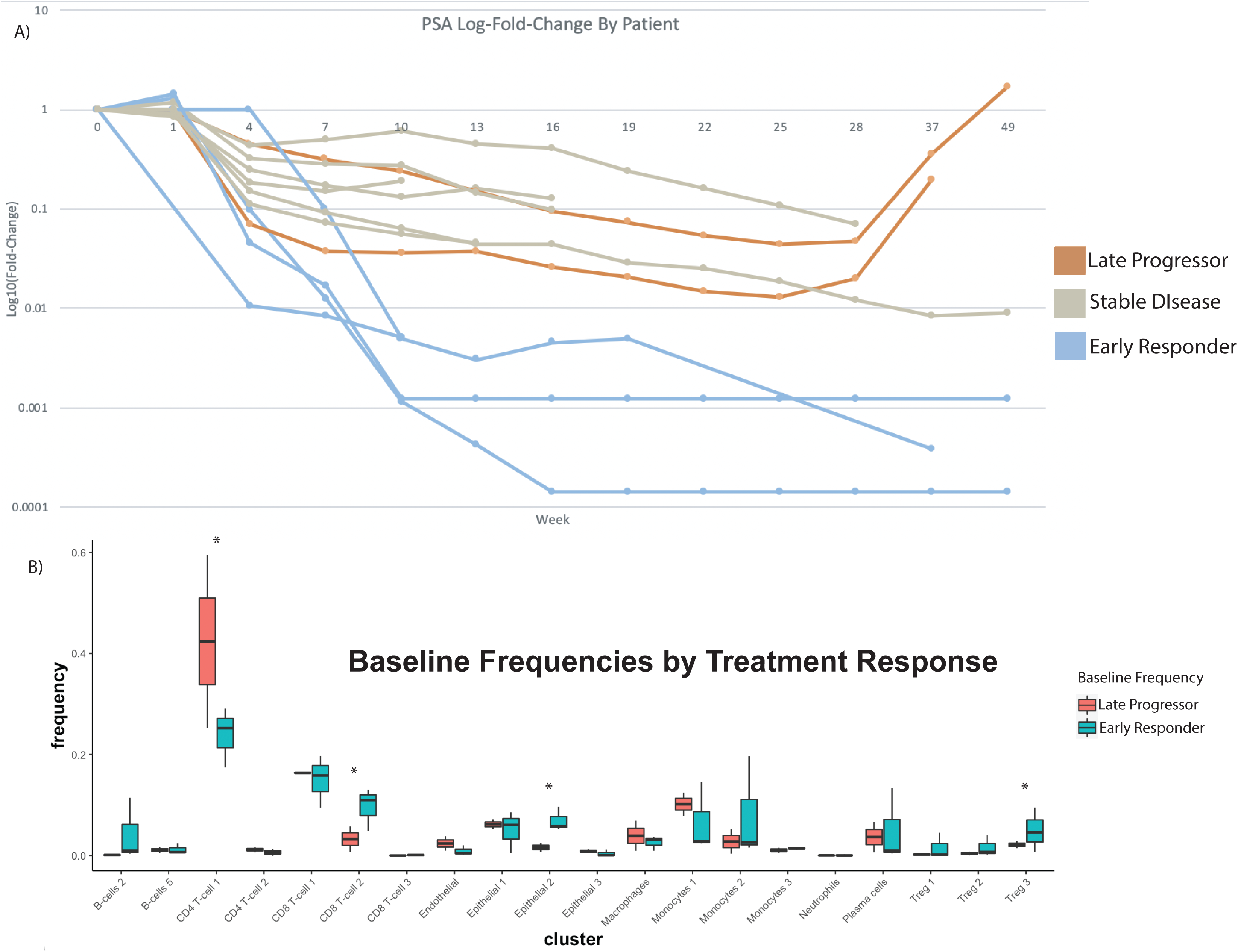
Differences in Baseline Immune Composition Associate with Differences in Treatment Response. A) Spider-plot of log10(Fold-Change) from Baseline in Prostate-Specific Antigen (PSA) over time with treatment, for each patient, such that four patients, labelled in blue, exhibited rapid and dramatic decrease to below 1% of initial PSA and were identified as Early Responders to treatment, and two patients, labelled in orange, initially responded to treatment with a rapid increase in PSA observed after on-treatment week 28. These were considered Late Progressors on-treatment. The remaining patients, in grey, generally trended toward a decreasing PSA, though not as rapidly as the Early Responders. B) Boxplot showing distribution of frequencies at Baseline for each cell cluster, comparing frequencies in Early Responders vs Late Progressors, such that clusters with significant difference at baseline (p<0.05 by Student’s T-test) included CD4 T-cell 1, CD8 T-cell 2, Treg 3, and Epithelial 2.

Overall, at baseline, higher Cluster 2 LAG3+ CD8 T cells representation was significantly associated with early PSA responders (*p* = 0.007). Interestingly, this represents the same population of LAG3+ CD8 T-cells whose representation was expanded following anti-PD1 therapy. The specific subpopulation of TNFRSF18+ T regulatory cells (Cluster 3 Tregs) also trended with an early PSA response, even though the association was not statistically significant (*p* = 0.14). Conversely, over-representation of Cluster 1 CD4 T cells was significantly associated with late PSA progression (*p* = 0.026). Notably, one of the most differentially active proteins in this CD4 subtype was tumor necrosis family (TNF), a multifunction proinflammatory cytokine implicated in tumor progression ^55–57^.

### Transformed Cells Show Transcriptional Heterogeneity Across Metastatic Sites

Initial analysis of tumor cells by protein activity-based clustering resulted in three ‘epithelial’ clusters (EPI_1_, EPI_2_, and EPI_3_) (Figure 1B). Copy number alteration (CNA) inference and expression of the KLK3 prostate tumor marker gene confirmed that all three subtypes represent malignantly transformed epithelial cells (Figure S6). Following identification of transformed epithelial cells across pre-treatment samples from all metastatic sites, we found that all three epithelial subtypes were represented in bone, lymph node, and lung-derived samples. To further refine the heterogeneity of the tumor cell compartment, we performed a more stringent cluster analysis after excluding all non-transformed cells. The analysis yielded eight molecularly distinct, yet co-existing tumor cell subtypes (REF-EPI_1_ – REF-EPI_8_) (Figure 5A) that were further analyzed to assess enrichment of Cancer Hallmarks and cancer-related pathways in differentially active proteins (Figure S5). Intriguingly, REF-EPI_1_ was the subtype most enriched in androgen response proteins, REF-EPI_2_ and REF-EPI_3_ were defined by upregulation of E2F and MYC targets, and G2M checkpoint proteins, REF-EPI_4_ and REF-EPI_5_ were defined by upregulation of TNFα signaling and interferon response proteins, REF-EPI_6_ was identified by heme metabolism, REF-EPI_7_ by unfolded protein response and androgen response, and finally REF-EPI_8_ by activation of the reactive oxygen species pathway. The most differentially upregulated proteins in each cluster are shown in Figure 6.

**Figure 5:**
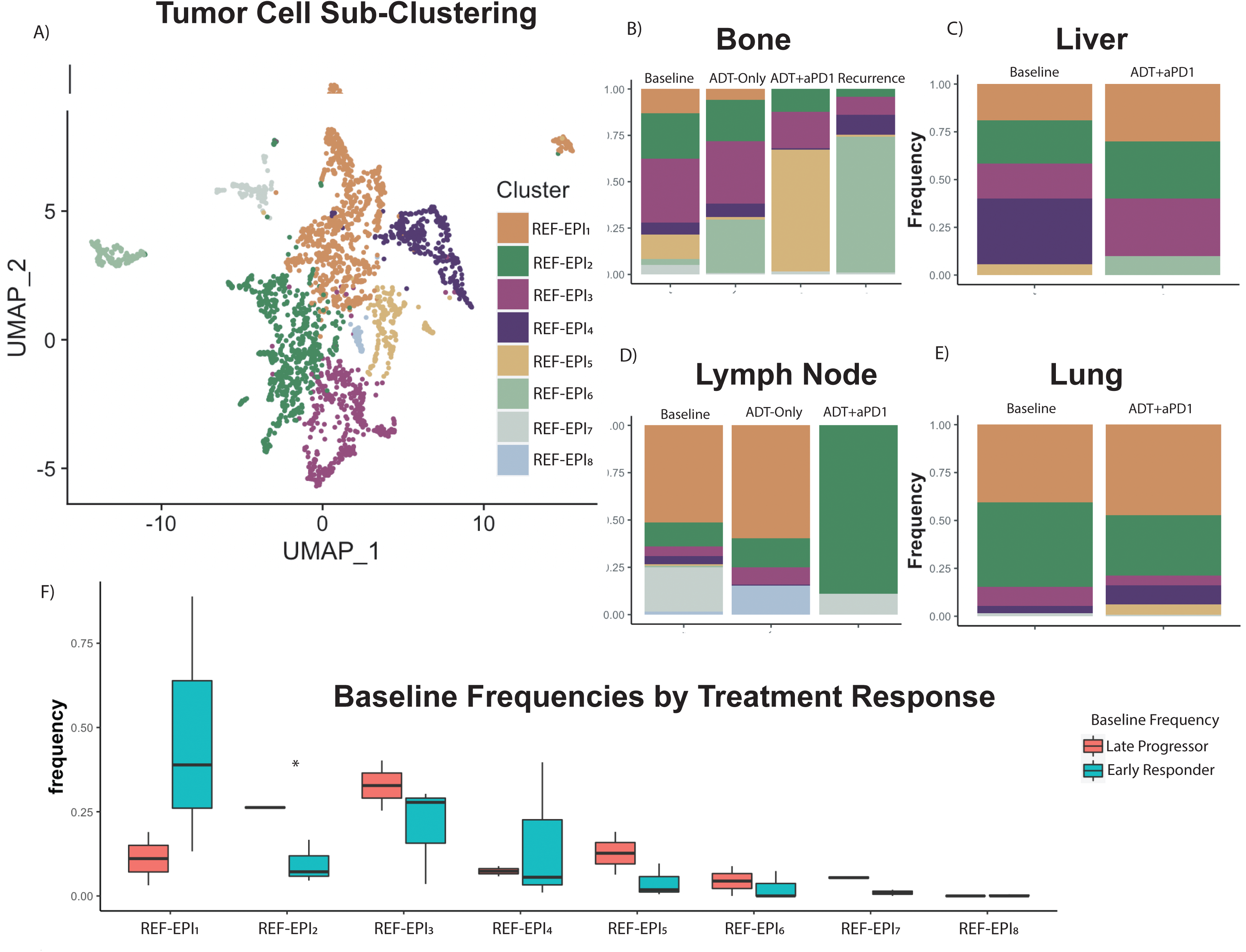
Sub-Clustering Reveals Heterogeneity of Tumor Cells by Tissue Site. A) UMAP plot showing sub-clustering by resolution-optimized Louvain algorithm of only tumor cells (Epithelial 1, Epithelial 2, and Epithelial 3 from Figure 1B). Plot shows aggregate of all 2,550 tumor cells across all patients at all time-points. B) Stacked barplot of tumor cluster frequency by treatment time-point in patients with metastatic Bone tumors. C) Stacked barplot of tumor cluster frequency by treatment time-point in patients with metastatic Liver tumors. D) Stacked barplot of tumor cluster frequency by treatment time-point in patients with metastatic Lymph Node tumors. E) Stacked barplot of tumor cluster frequency by treatment time-point in patients with metastatic Lung tumors. F) Boxplot showing distribution of frequencies at Baseline for each tumor subcluster, comparing frequencies in Early Responders vs Late Progressors, such that the only cluster with significant difference at baseline (p<0.05 by Student’s T-test) was REF-EPI_2_, with higher baseline frequency in Late Progressors.

**Figure 6:**
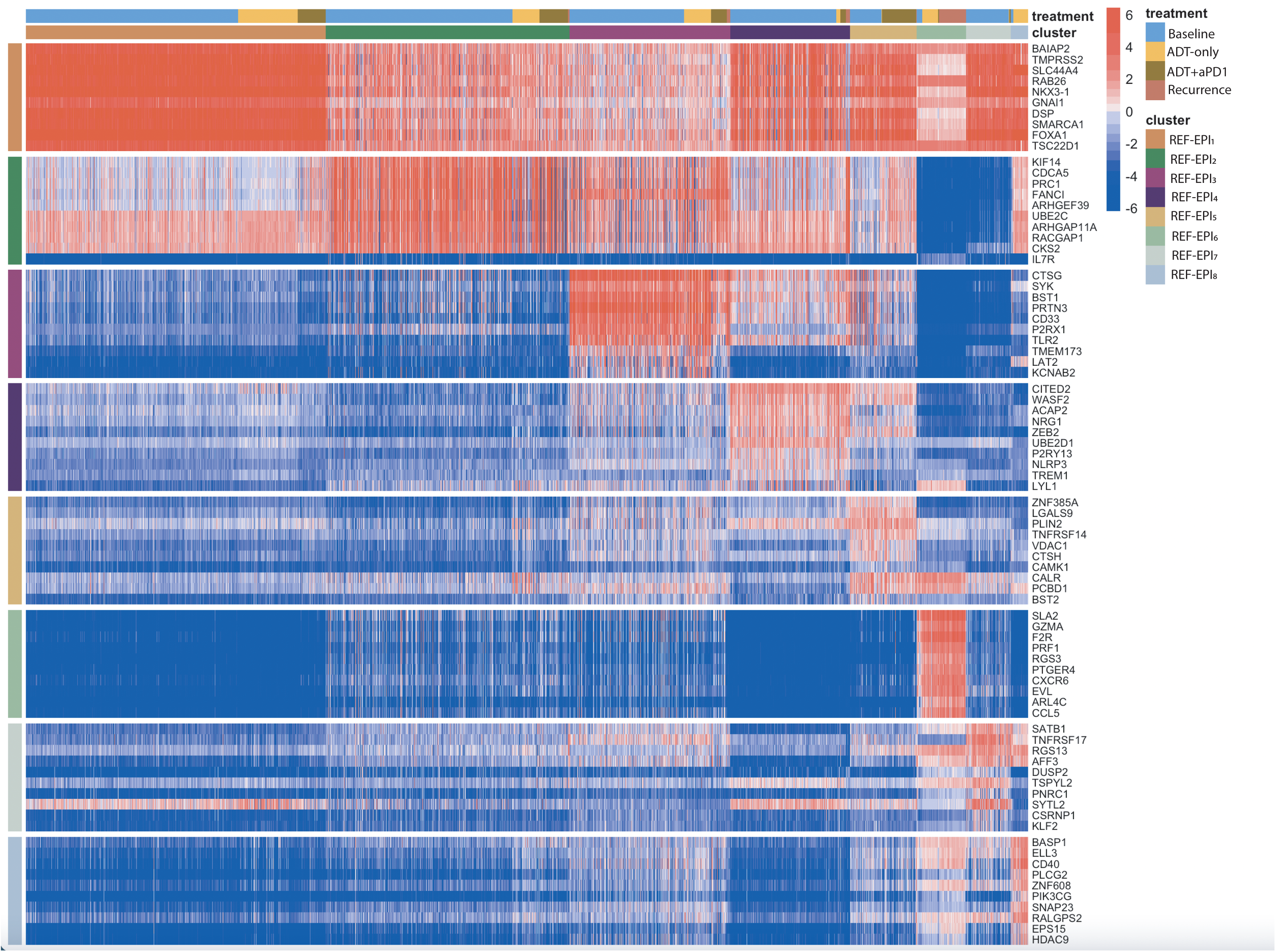
Top Protein Activity Tumor Subcluster Markers. Heatmap of top 10 most differentially activated proteins for each tumor subcluster in Figure 5 from aggregate single-cell RNA-Sequencing data across all patient samples. Each row represents a protein, grouped by cluster in which they are the most active, with cluster labels on the x and y-axes. Each column represents a single cell. Above the x-axis cluster label there is also a treatment label indicating timepoint at which a given cell was profiled.

Stacked frequency bar plots (Figure 5B-E) show changes in the relative fraction of each transformed tumor cell subtype, across different metastatic sites at baseline, after ADT treatment, after combination therapy, and at recurrence, in available biopsy samples. At baseline, there was wide variability in the composition of tumor subclusters across the metastatic sites. Bone and lymph node-derived samples were more heterogenous, with representation of nearly all tumor cell subtypes. In contrast, liver and lung-derived samples presented lower heterogeneity. In particular, lung was comprised almost entirely of cells from the REF-EPI_1_ and REF-EPI_2_ subtypes. This is interesting because patients with pulmonary-only prostate cancer metastases appear to have favorable overall prognosis ^58^. After treatment, tumor subtype representation was differentially affected across distinct metastatic sites. In bone samples, we observed relative increase of REF-EPI_6_ cells following ADT treatment (*p* = 4.4e-14). With the addition of anti-PD-1 REF-EPI_6_ cells became almost undetectable (*p* = 0.02), while REF-EPI_5_ cells increased to comprise 65% of tumor cells (*p* = 9.9e=27). However, in the recurrent bone sample, REF-EPI_6_ cells comprised nearly 75% of all tumor cells, while REF-EPI_5_ were reduced to only 1% of all tumor cells (Figure 5B), suggesting that REF-EPI_6_ may play an important role in drug resistance.

In lymph node-derived, pre-treatment samples, REF-EPI_1_ cells accounted for nearly 50% of all tumor cells. Yet, after ADT, their representation increased to ∼65%. Similar to bone-derived samples, following ADT treatment, the fractional representation of tumor cell subtypes in lymph node-derived samples changed slightly, while preserving the original heterogeneity. However, addition of anti-PD-1 therapy led to emergence of REF-EPI_2_ as a predominant subtype, comprising 89% of the cells (*p* = 9.4e-7). Liver and lung-derived samples also maintained heterogeneity but did not show emergence of a predominant tumor cell subtype after combination therapy. As discussed, there were no patients with liver and lung metastases randomized to the ADT only timepoint.

We next analyzed whether the presence of individual tumor cell subtypes at baseline was associated with differential PSA response (early responders vs. late progressors). Indeed, REF-EPI_1_, associated with an androgen response signature, was significantly overrepresented in baseline samples from ‘early responders’ (*p* = 0.05) compared to ‘late progressors.’ In contrast, REF-EPI_2_ and REF-EPI_3_ were overrepresented in baseline samples from ‘late progressors’, who ultimately failed to respond to treatment (*p* = 0.0008 and 0.08, respectively) (Figure 5F).

Interestingly, two proteins (TMPRSS2 and NKX3-1), which are regulated by the androgen receptor (AR), were among the most differentially active in REF-EPI_1_ ^59–61^. In contrast, there were no AR-regulated proteins in among the most differentially active in other tumor cell subtypes. Taken together, these data imply that activation of AR-regulated proteins in subpopulations representing the majority of tumor cells is associated with early treatment response. Notably, REF-EPI_2_ is characterized by aberrant activity of KIF14 (most-upregulated protein), which has previously been described as a candidate oncogene correlating with poor prognosis in prostate cancer ^49^.

### Identification of candidate targets in molecularly distinct subtypes of tumor cells

To identify candidate druggable proteins aberrantly activated in each of the 8 tumor cell subtypes, we leveraged the OncoTarget algorithm ^28,29^ (Figure 7). Most significantly, REF-EPI_2_ and REF-EPI_3_, which are associated with tumor progression on treatment, lacked activity of druggable proteins shared by other subtypes; most notably, no activity of the androgen receptor protein (AR) was detected. AR activity was highest in REF-EPI_1_, which was most enriched in early responders. However, the two poor-prognosis subtypes, REF-EPI_2_ and REF-EPI_3_, shared elevated activity of other druggable proteins, such as topoisomerase 2-alpha (TOP2A), for which a number of FDA-approved and investigational compounds are annotated in DrugBank ^62^, including Doxorubicin and Etoposide. REF-EPI_3_ also presented uniquely elevated activity of CD33, a surface receptor representing a high-affinity target of the FDA approved drug Gemtuzumab ozogamicin and of the investigational agent AVE9633.

**Figure 7:**
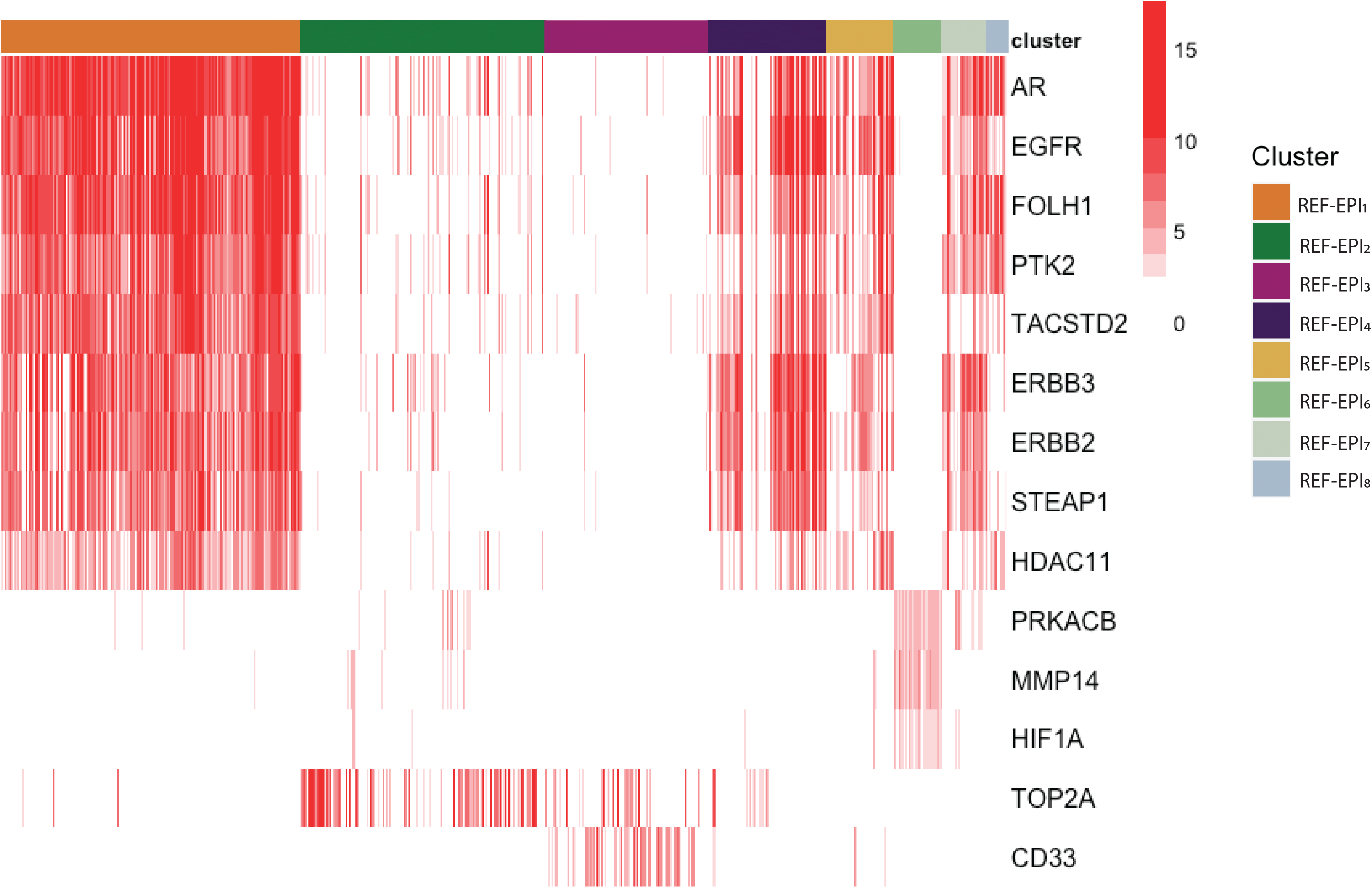
OncoTarget Druggable Proteins in Each Tumor Cell Subcluster. Single-Cell heatmap of all druggable proteins from DrugBank, active with median -log10(p-value) > 5 in any tumor cell subcluster, as inferred by OncoTarget. REF-EPI_2_ and REF-EPI_3_ are the most phenotypically distinct with respect to druggable protein activity, as they do not have high activity of AR and are characterized instead by high activity of TOP2A. REF-EPI_6_, which was specifically enriched in post-treatment recurrence for the Late Progressor sample with post-recurrence single-cell RNA-Seq, is also phenotypically distinct, with activity of PRKACB, MMP14, and HIF1A.

Other druggable proteins with high activity in REF-EPI_1_, as well as in REF-EPI_4_, REF-EPI_5_, REF-EPI_7_, and REF-EPI_8_ include EGFR, FOLH1, PTK2, TACSTD2, ERBB2, ERBB3, STEAP1, and HDAC11 (Figure 7). Finally, REF-EPI_6_, which was expanded in the bone metastasis of the patient profiled at time of recurrence (Figure 5B), also presented a unique druggable protein profile, characterized by low activity of AR and of other druggable proteins seen in REF-EPI_1_, REF-EPI_4_, REF-EPI_5_, REF-EPI_7_, and REF-EPI_8_ yet high activity of druggable proteins PRKACB, MMP14, and HIF1A. This represents a profile for which frequency may need to be assessed in additional patients with disease recurrence. It may represent a rare and more aggressive prostate tumor cell phenotype, even though it did not associate with differences in treatment response across patients at baseline (Figure 5F). Rather it was represented exclusively in a single outlier patient.

### Validation of Association Between Tumor Cell Clusters and Outcome

To assess the generalizability of our findings with respect to tumor cell subcluster associations with treatment response, we defined each set of differentially active proteins in each subcluster as a unique protein activity signature for that cluster (Figure 6). With these signatures and our protein activity inference algorithm, we were able to test enrichment of each cluster within larger cohorts of bulk-RNA-Sequencing data. As there are no previously published cohorts of metastatic prostate cancer patients treated with combination ADT plus anti-PD1 immunotherapy, we assessed the general prognostic significance of each tumor cell population across treatments in the TCGA dataset. By Cox regression on patient-by-patient normalized enrichment scores (Figure 8A), enrichment of tumor cell REF-EPI_2_ was significantly associated with shorter recurrence-free-survival (hazard ratio 1.37, p = 0.002). The leading-edge genes in the REF-EPI_2_ protein activity signature most enriched in patients with recurrence compared to non-recurrence are reported in Figure 8B and include KIF14 as well as TOP2A, both determined to biologically significant markers of REF-EPI_2_.

**Figure 8:**
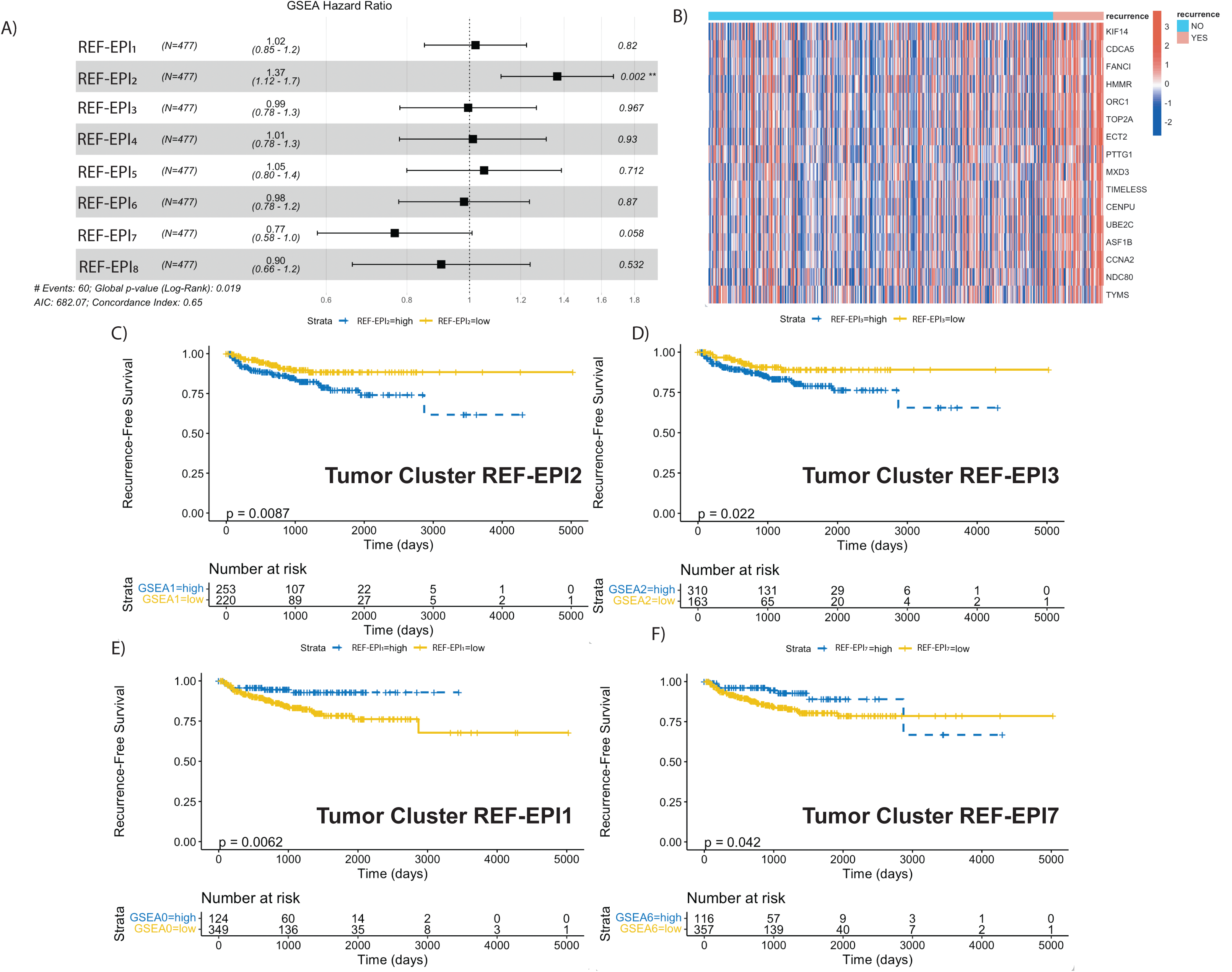
Tumor Single-Cell Subcluster Signatures Associate with Differential Outcomes in TCGA. A) Forest plot of Cox regression hazard ratios testing association in TCGA of patient-by-patient Normalized Enrichment Score for each tumor subcluster gene set with Recurrence-Free survival. REF-EPI_2_ gene set enrichment is significantly associated with worse survival outcomes (p = 0.002). B) Heatmap of Leading-Edge Gene Set from REF-EPI_2_ comparing all Recurrent vs Non-Recurrent patients in TCGA. C) Kaplan-Meier curve testing association of binarized REF-EPI_2_ gene set enrichment (greater than 0 = high, less than 0 = low) with recurrence-free survival in TCGA, such that REF-EPI_2_ enrichment significantly associates with worse recurrence-free survival. D) Kaplan-Meier curve testing association of binarized REF-EPI_3_ gene set enrichment (greater than 0 = high, less than 0 = low) with recurrence-free survival in TCGA, such that REF-EPI_3_ enrichment significantly associates with worse recurrence-free survival. E) Kaplan-Meier curve testing association of binarized REF-EPI_1_ gene set enrichment (greater than 0 = high, less than 0 = low) with recurrence-free survival in TCGA, such that cluster 0 enrichment significantly associates with improved recurrence-free survival. F) Kaplan-Meier curve testing association of binarized REF-EPI_7_ gene set enrichment (greater than 0 = high, less than 0 = low) with recurrence-free survival in TCGA, such that REF-EPI_7_ enrichment significantly associates with improved recurrence-free survival, up to 2800 days. Kaplan-Meier curves are not shown for the remaining clusters as log-rank p-values for these were not statistically significant (p>0.05).

Log-rank testing of enrichment scores binarized to “high” vs “low” showed significant association of both REF-EPI_2_ and REF-EPI_3_ protein activity profiles with shorter recurrence-free survival (*p* = 0.0087 and 0.022, respectively) (Figure 8C-D), and significant association of REF-EPI_1_ and REF-EPI_7_ protein activity profiles with improved recurrence-free survival (p = 0.0062 and 0.042, respectively) (Figure 8E-F). No other subtype specific signature was statistically significantly associated with survival. In two smaller datasets specifically profiling metastatic CRPC tumors (East Coast Stand Up to Cancer, West Coast Stand Up to Cancer) ^39,40^, trends were observed toward association of REF-EPI_2_ and REF-EPI_3_ with worse overall survival (Figure S3, Figure S4). Specifically, REF-EPI_3_ was significantly associated with worse overall survival in East Coast SU2C (*p* = 0.045) (Figure S4B), while REF-EPI_2_ did not achieve statistical significance. However, neither dataset includes recurrence-free survival or PSA response as clinical metadata. Taken together, these results are highly concordant with the concept that overrepresentation of REF-EPI_1_ cells at baseline associate with better treatment response while overrepresentation of REF-EPI_2_ and REF-EPI_3_ at baseline associate with worse treatment response.

## DISCUSSION

Evaluation of primary prostate cancer and metastatic, <u>castration-resistant</u> disease using high-throughput, transcriptomic sequencing,^23,52,63–65^ showed that the TME is relatively immune-depleted. Here, we used scRNA-seq profiles to comprehensively characterize the TME of metastatic, <u>castration-sensitive</u> prostate cancer (mCSPC), across a variety of tissue types. Using longitudinal samples from 10 patients over a treatment course with ADT and anti-PD-1 antibody, we describe the baseline TME and tumor cells, the specific changes induced with treatment, and associated baseline features with PSA response. We leveraged our expertise in inferred protein-activity computational methods to increase resolution of the immune and tumor cell subpopulations as compared to conventional gene-expression and transcriptomic methods. In particular, protein activity enabled stratification of tumor cell subpopulations with significantly different drug targets and variable associations with post-treatment outcome, whereas tumor cells did not sub-cluster by gene expression at all due to excessive transcriptional background noise.

Profiling transcriptomes from a cumulative 40,270 single-cells, our study uncovered a previously unsuspected rich immune infiltrate in untreated mCSPC samples. In our analyses the baseline bone, lymph node, and liver samples were similarly immune infiltrated while the lung metastasis was relatively immune-depleted (Figure 1C,D) and distinctly different. The latter should be considered with the caveat of a small sample size. These data add to the notion that pulmonary-tropic and non-pulmonary metastatic mCSPC may be biologically and/or immunologically fundamentally different ^58^. Regarding malignantly transformed epithelial cells in mCSPC samples, our data adds to the body of literature by demonstrating both intra- and inter-patient tumor cell heterogeneity ^66^ . However, it significantly expands our knowledge by identifying 8 distinct tumor cell subpopulations and reporting their different representation in patients with differential response to treatment and in pre- vs. on-treatment samples (Figure 5A) (Figure 5B-E). These subpopulations were invisible by gene expression profiling and could only be detected by VIPER-based protein activity analysis. Specifically, we show phenotypic changes in tumor cell types induced by treatment, associate baseline tumor cell phenotypes with clinical response, and define pathways enriched longitudinally and upon recurrent, progressive disease.

Our study is limited with respect to the total number of patient samples per tissue type and analysis of a single metastatic site per patient. However, sample size was sufficient to assess statistical significance based on over or under representation of specific subpopulations in the mCSPC TME. In addition, longitudinal analysis of prostate cancer tumor metastases at the single cell level, over a course of treatment, has not been previously reported. Our analyses are potentially biased towards more aggressive biology given that only patients with evaluable disease at the time of on-treatment biopsy were able to safely undergo metastatic biopsy. As such, we are unable to comment regarding the on-treatment changes in the TME and tumor cell profiles of participants who were rapidly responding to therapy. The time points for on-treatment biopsies are fixed due to the nature of a clinical trial and are not based on tumor kinetics although this may be an option in future studies. Here we comment on cell types that were represented across all tissue types to avoid analyzing subpopulations that may be less relevant to tumor-immune crosstalk given their expected presence in a specific metastatic niche, *e.g.,* common progenitor cells in the bone marrow and B cells in the lymph nodes. This approach highlights broad changes in transcriptional program across tissue types in lieu of a deep dive on tissue specific idiosyncrasies.

The observation that treatment induces changes in both the TME cellular composition and transcriptional program of tumor cells, so called lineage plasticity, is consistent with other studies in prostate cancer ^67–69^. However, the TME changes we observed with administration of ADT in mCSPC are opposite of those described in primary prostate cancer. In primary prostate cancer, an immune infiltrate rich in T cells invades the TME after ADT administration ^6,12^. In mCSPC biopsy samples we observed a decrease in CD4 and CD8 T cells after ADT, whereas the combination of ADT and anti-PD-1 immunotherapy was effective at recruiting CD8 effector T cells. It is possible that these observed differences between primary and metastatic CSPC are due to the baseline TME composition, *i.e.,* an ‘immune desert’ vs. ‘immune replete’ respectively, and immunomodulatory factors already present in the milieu.

Importantly, in our study, we observed significant increases in TNF-α+ CD4 non-T reg T cells (CD4 1), LAG3+ CD8 T cells (CD8 2), and GITR+ T regs (Treg 3) after combination therapy. This highlights the notion that combination therapy with ADT and anti-PD-1 therapy in men with mCSPC is an immunologically active combination, even in bone metastasis. Lymphocyte activation gene-3 (LAG3, CD223), a CD4 homologue that binds to MHC class II ^53^, is upregulated on CD8 T cells after antigen experience and represents an ‘exhausted’ state, which negatively regulates their activation and homeostasis ^70,71^. Dual-inhibition of PD-1 and LAG-3 was recently shown to improve outcomes in patients with melanoma in a large phase 3 clinical trial ^72^. In prostate cancer models that are resistant to single-agent PD-1, dual blockade of PD-1 and LAG3 improved vaccine efficacy providing evidence that combination immune checkpoint therapy may be key to improving clinical response outcomes ^73^. Taken together, these data suggest that LAG3 may be a potential adjunct to combination immune checkpoint therapy in prostate cancer. Glucocorticoid-induced TNFR-Related (GITR) protein, an immune checkpoint receptor, belongs to Tumor Necrosis Factor Receptor Superfamily (TNFRSF). GITR is preferentially expressed on CD8 and T reg cells and agonistic antibodies are shown to potentiate the former and reduce functionality of the latter ^74–76^. Although several preclinical and early phase studies have shown that anti-GITR agonist antibodies are safe, clinical results have been modest thus far ^77,78^. Trials of dual immune checkpoint blockade targeting GITR and PD-1 have shown slight advantage over single-agent anti-GITR agonists antibodies ^79,80^, although data in mCSPC is somewhat limited. Tumor Necrosis Factor alpha (TNF-α), a major inflammatory cytokine with signaling potential both as a membrane-bound protein and as a soluble ligand, was initially implicated as an anti-tumor cytokine but has since been connected, in complete contraindication to its name, to tumor progression ^55,56,81^. Given the ever-increasing number of patients treated with combination immune checkpoint therapy, more patients are developing immune-related adverse events (irAEs) that frequently require treatment with immunosuppressive therapies like anti-TNF-α inhibitors. As such, much has been learned about the effects of anti-TNF-α inhibitors in patients with cancer ^56^. Several studies that aimed to abrogate irAEs prior to their onset used TNF-α inhibitors upfront with combination immune checkpoint blockade and yielded improvements in anti-tumor efficacy ^57^ suggesting a role for TNF-α blockade as an anti-tumor agent. Our group has shown that elevated TNF-α levels are associated with PSA progression in men with biochemically recurrent prostate cancer, consistent with other studies in metastatic prostate cancer ^82–84^. Others have shown that TNF-α promotes prostate cancer dissemination from metastatic lymph nodes through activation of the CCL21/CCR7 axis ^85^ and that inhibiting TNF-α, and its downstream prosurvival signaling molecules such as NF-κB and Bcl-2, is a potent therapeutic agent for androgen-independent prostate cancer ^86^. Taken together, these data in the context of increased protein activity of TNF-α in CD4 T cells after ADT and anti-PD-1 therapy shown here (Figure 2, S5), suggest that inhibiting TNF-α, concurrently or sequentially, with ADT and immune checkpoint therapy in CSPC may be an effective treatment combination.

Due to the tropism of metastatic prostate cancer to bone, tissue-specific changes occurring in the TME of bone samples were of particular interest. Notably, following ADT alone, we observed a relative increase of myeloid cells (Figure 3A-B) and a noticeable decrease in CD4 non-T reg T cells as well as tumor cells. As discussed earlier, the observed expansion of CD8 T cells was most pronounced in the bone TME after combination therapy compared to other tissue types (Figure 3B). This highlights the notion that moving away from a ‘one-therapy-treats-all’ treatment paradigm and towards more precision-based, targeted, and tissue-based algorithms is likely on the horizon.

Given that our transcriptomic data comes from a prospective clinical trial with close monitoring of clinical response, associations between baseline biopsy features and PSA response was possible. Thus far, the primary endpoint of our phase 2 trial, the rate of undetectable PSA (< 0.02 ng/dL) at 37 weeks after combination therapy, is 42%. This compared to 32% in the ADT plus docetaxel arm of the phase 3 CHAARTED trial (E3805) ^87^. Although the proportional difference in the rates of undetectable PSA between our small phase 2 study and CHAARTED are not currently statistically significant, due to current statistical power and enrollment, we remain encouraged. It is somewhat perplexing that the rate of undetectable PSA is not overwhelmingly lower given the robust influx of CD8 effector T cells we observed with combination therapy.

New observations from our data sheds light on issues using classically defined immune markers as predictive biomarkers. As previously noted, the association of baseline immune subclusters with PSA response showed that TNF-α+ CD4 T cells (CD4 1) were statistically significantly associated with late PSA progression, and also that LAG3+ CD8 T cells (CD8 2) and GITR+ T reg cells (T reg 3) were both statistically significantly associated with early PSA response. In the case of T regs this is somewhat paradoxical. T regulatory cells are classically considered to be an immune suppressive subpopulation of T cells. However, a recent analysis of Tregs using high-dimensional flow cytometry from an NPK-C1 transplantable prostate tumor model revealed significant phenotypic diversity within Tregs, including a Treg subpopulation enriched in regressing tumors ^88^. Thus, our data supports this preclinical observation that ‘favorable’ Tregs may be present in the TME at various states of treatment pressure. Further, taken together, these data demonstrate the capacity of single-cell and high-dimensional data to provide more granularity on immune subpopulations and may even challenge the definitions of classical ‘immunosuppressive’ or ‘immune effector cells.’

We observed both expected and unexpected findings upon association of baseline tumor subclusters with PSA response. REF-EPI_1_, which has high protein activity levels of two androgen receptor (AR) regulated proteins (TMPRSS2 and NKX3-2), was, not surprisingly, associated with early PSA response (Figure 5F). Using the ‘hallmarks of cancer’ pathway enrichment analysis ^41^, we confirmed that REF-EPI_1_ was indeed enriched with androgen response genes (Figure S5). We confirmed a positive association between this gene set and clinical outcomes in the TCGA dataset using GSEA (Figure 8E). In contrast, REF-EPI_2_ and REF-EPI_3_, defined by upregulation of E2F targets, Myc targets, and G2M checkpoint (Figure S5), were associated with late PSA progression (Figure 5F). These associations were validated in a similar manner (Figures 8C-D, S4). Interestingly, we did not observe tumor REF-EPI_4_, the subcluster enriched with TNF-α signaling genes (Figure S5), to be associated with worse outcomes (Figure 5F). This association is opposite to what we observed in the TNF-α+ CD4 T cells (CD4 1), which was associated with decreased clinical outcomes. Taken together, these observations highlight the notion that the specific cell of protein activity and/or expression (CD4 T cells vs. tumor cells) indeed matters and is a reflection of the underlying immunologic processes at play.

Because certain tumor subclusters were associated with PSA or metastatic progression, we used OncoTarget (see methods) to examine potential druggable proteins active in the tumor subclusters. REF-EPI_2_ and REF-EPI_3_ showed high activity of TOP2A and low AR activity suggesting these subclusters may be upfront resistant to AR-targeted therapies (Figure 7). Although REF-EPI_2_ and REF-EPI_3_ were associated with PSA progression in our dataset (Figure 5F), and validated as such in external datasets, they were not the dominant tumor subclusters seen in the tumor progression biopsy (Figure 5B). More metastatic biopsies at the time of tumor recurrence would be helpful to delineate if there is a role for targeting tumor REF-EPI_2_ and REF-EPI_3_ with topoisomerase inhibition in mCSPC. Targeting these tumor cell populations upfront to eradicate them prior to combination ADT and anti-PD-1 therapy may be one approach for future clinical trials. Of particular interest was tumor cluster REF-EPI_6_, the dominant cluster seen in the bone progression biopsy. Figure 7 shows that potential druggable proteins in REF-EPI_6_ include PRKACB (cAMP-dependent protein kinase catalytic subunit beta), MMP14 (matrix metallopeptidase 14), and HIF1a (Hypoxia Inducible Factor 1 Subunit Alpha). Taken together, our combined dataset and analyses highlight several potential targets for worthy of further investigation in mCSPC.

Building on this rich longitudinal transcriptomic dataset, we propose that the “holy grail” treatment for men with mCSPC will require a multi-pronged and adaptive combination regimen to elicit complete and durable responses. We propose a term ‘Highly Active Anti-Tumor Therapy’ (or HAATT), that includes a treatment regimen with the following properties: 1) strong upfront tumor cell killing perhaps directed at known resistant tumor cell clones, 2) activation of CD8 effector T cell function via combination immune checkpoint therapy (anti-PD-1 with anti-LAG3, anti-GITR agonist antibodies, or novel agents targeting costimulatory agonists like 4-1BB or B7-H3), 3) depletion or blocking of T regulatory cells, and 4) targeting immunosuppressive or tumor-permissive molecules in the TME (*i.e.*, cytokines, chemokine receptors, metabolomic pathways, or transcription factors). As demonstrated, it is imperative to review the underlying tumor immunology and biology continuously and critically with advanced techniques when conceiving of the next biologically plausible clinical trials.

## Acknowledgements

This work was supported by grants from the Conquer Cancer Foundation (2019YIA to JEH), the Prostate Cancer Foundation (2020YIA to JEH) and the National Institutes of Health (T32CA203703 to JEH, UL1TR001873 to JEH, F30CA260765-01 to AZO), as well as by the 54CA209997, 35CA197745, S10OD012351 and S10OD0217640 to AC. We also acknowledge the P30CA013696 Cancer Center grant for the support of the Cancer Biostatistics Shared Resource and JP Sulzberger Columbia Genome Center at the Herbert Irving Comprehensive Cancer Center. Regeneron Pharmaceuticals provided the funding for the clinical trial.

## Figure Captions

**Figure S1: Gene Expression Clustering**

A) UMAP plot showing clustering of all cells in tumor microenvironment across all patients, clustered on gene expression instead of VIPER-inferred protein activity. Cell types are inferred by SingleR. Total number of clusters is smaller than clustering in Figure 1 on VIPER-inferred protein activity. B) UMAP plot from A, split by metastatic tissue site.

**Figure S2: Top Gene Expression Cluster Markers**

Heatmap of top 5 most differentially upregulated genes for each cell type cluster from aggregate single-cell RNA-Sequencing data across all patient samples, with clusters corresponding to Figure S1. Each row represents a protein, grouped by cluster in which they are the most active, with cluster labels on the x and y-axes.

**Figure S3: Tumor Single-Cell Subcluster Signatures and Outcome in West Coast SU2C**

A) Forest plot of Cox regression hazard ratios testing association in West Coast Stand Up to Cancer (SU2C) dataset of patient-by-patient Normalized Enrichment Score for each tumor subcluster gene set with overall survival. B) Kaplan-Meier curve testing association of binarized REF-EPI_7_ gene set enrichment (greater than 0 = high, less than 0 = low) with survival, such that REF-EPI_7_ enrichment significantly associates with improved survival. Kaplan-Meier curves are not shown for the remaining clusters as log-rank p-values for these were not statistically significant (p>0.05).

**Figure S4: Tumor Single-Cell Subcluster Signatures and Outcome in West Coast SU2C**

A) Forest plot of Cox regression hazard ratios testing association in East Coast Stand Up to Cancer (SU2C) dataset of patient-by-patient Normalized Enrichment Score for each tumor subcluster gene set with overall survival. B) Kaplan-Meier curve testing association of binarized REF-EPI_3_ gene set enrichment (greater than 0 = high, less than 0 = low) with survival, such that REF-EPI_3_ enrichment significantly associates with worse survival. C) Kaplan-Meier curve testing association of binarized REF-EPI_7_ gene set enrichment (greater than 0 = high, less than 0 = low) with survival, such that REF-EPI_7_ enrichment significantly associates with improved survival. D) Kaplan-Meier curve testing association of binarized REF-EPI_8_ gene set enrichment (greater than 0 = high, less than 0 = low) with survival, such that REF-EPI_8_ enrichment significantly associates with improved survival. Kaplan-Meier curves are not shown for the remaining clusters as log-rank p-values for these were not statistically significant (p>0.05).

**Figure S5: Hallmarks of Cancer Enriched Pathways in Tumor Cell Subclusters**

For each tumor cell subcluster identified in Figure 5, plots of the top 10 enriched pathways from Hallmarks of Cancer. Pathway enrichment is computed on genes differentially expressed in each tumor subcluster relative to other tumor subclusters. -Log10(p-values) are plotted on the x-axes, such that statistically significant enriched pathways are shaded in blue.

**Figure S6: Identification of Tumor Cells by Marker Expression and Copy Number Variation**

A) log10 normalized expression of prostate cancer tumor marker protein KLK3 in each cell cluster identified by VIPER, such that expression is non-zero only in Epithelial cell clusters. B) InferCNV plot of cell-by-cell copy number variations, where all immune-lineage cells are taken as a copy-number-normal reference for inference of variations in copy number in Epithelial cell clusters and Endothelial cell cluster as a control. Each epithelial cell cluster is notable copy number aberrant across multiple chromosomes, while endothelial cells are grossly copy number normal.

## Notes

### Competing Interest Statement

Dr. Hawley has served as a paid consultant to Seagen and has received sponsored research funding to her institution from Regeneron and Dendreon. Dr. Drake is a co-inventor on patents licensed from JHU to BMS and Janssen and is currently an employee of Janssen Research. Dr. Califano is founder, equity holder, and consultant of DarwinHealth Inc., a company that has licensed some of the algorithms used in this manuscript from Columbia University. Columbia University is also an equity holder in DarwinHealth Inc. Dr. Lowy is an employee and stockholder of Regeneron Pharmaceuticals.

