## Supplemental Figures for "Single-Cell RNAseq Analysis Reveals Robust, Anti-PD-1-Mediated Increase of Immune Infiltrate in Metastatic Castration-Sensitive Prostate Cancer"

Figure S1

### Gene Expression UMAP

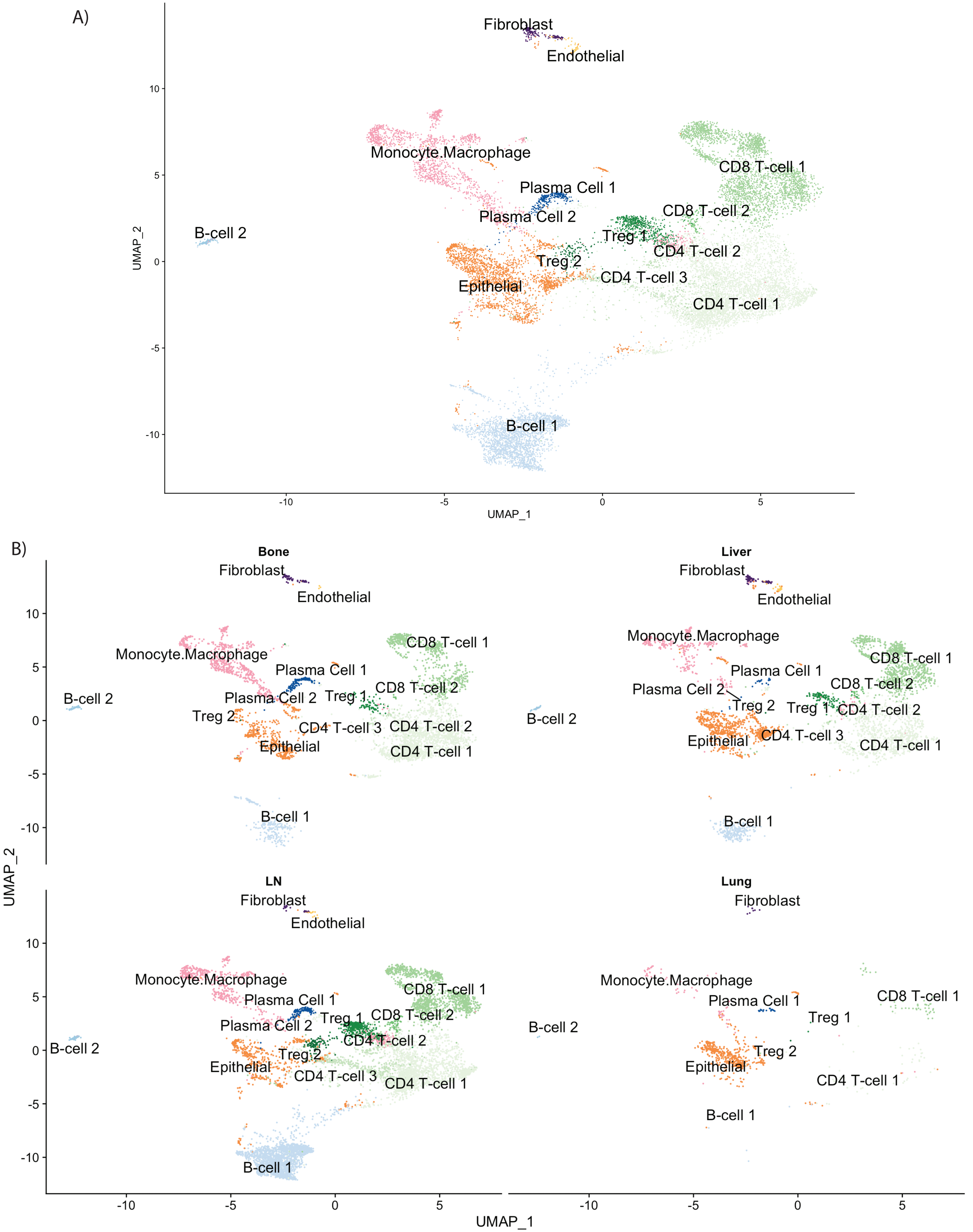

Figure S2

### Gene Expression Heatmap

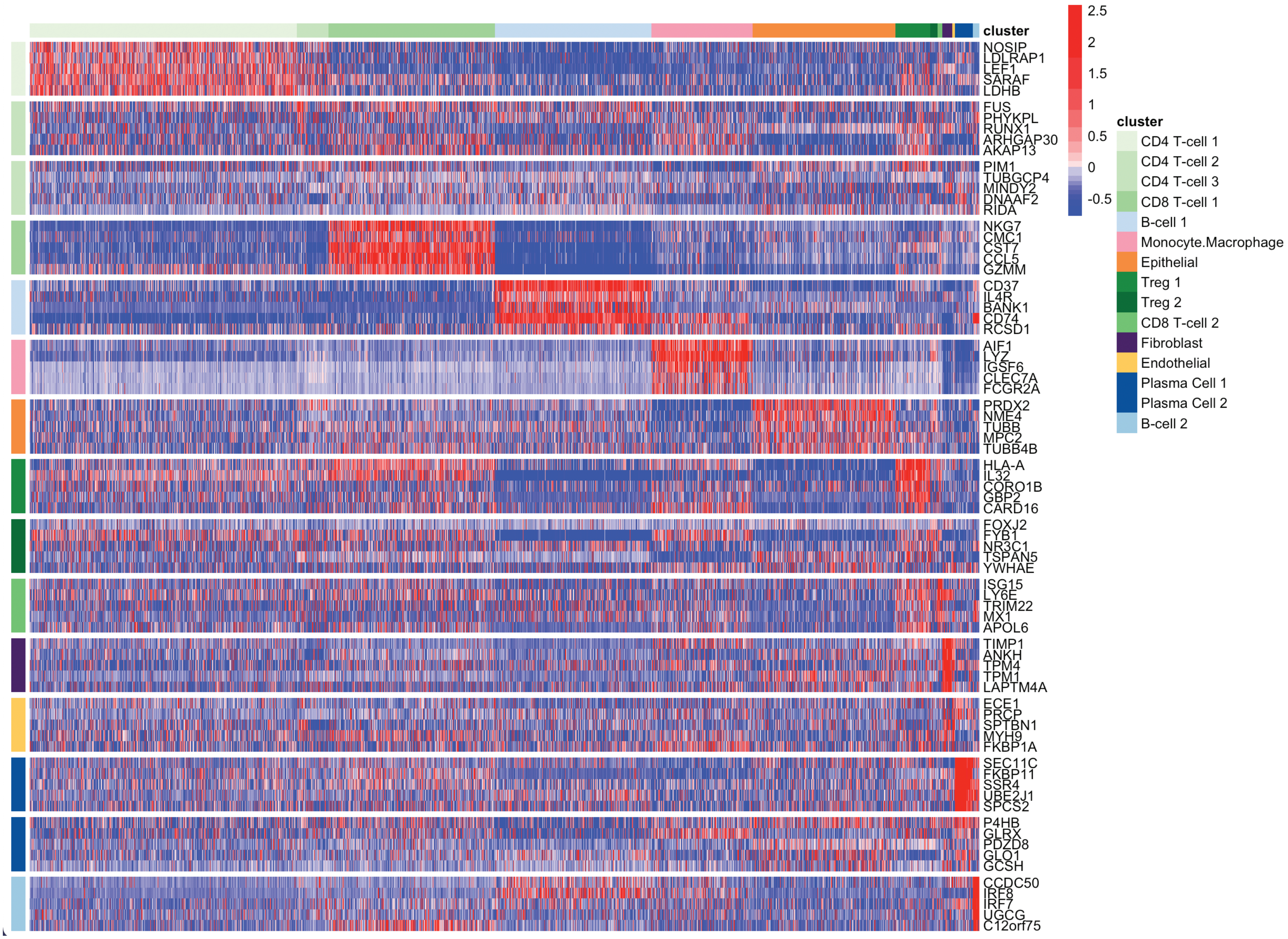

Figure S3

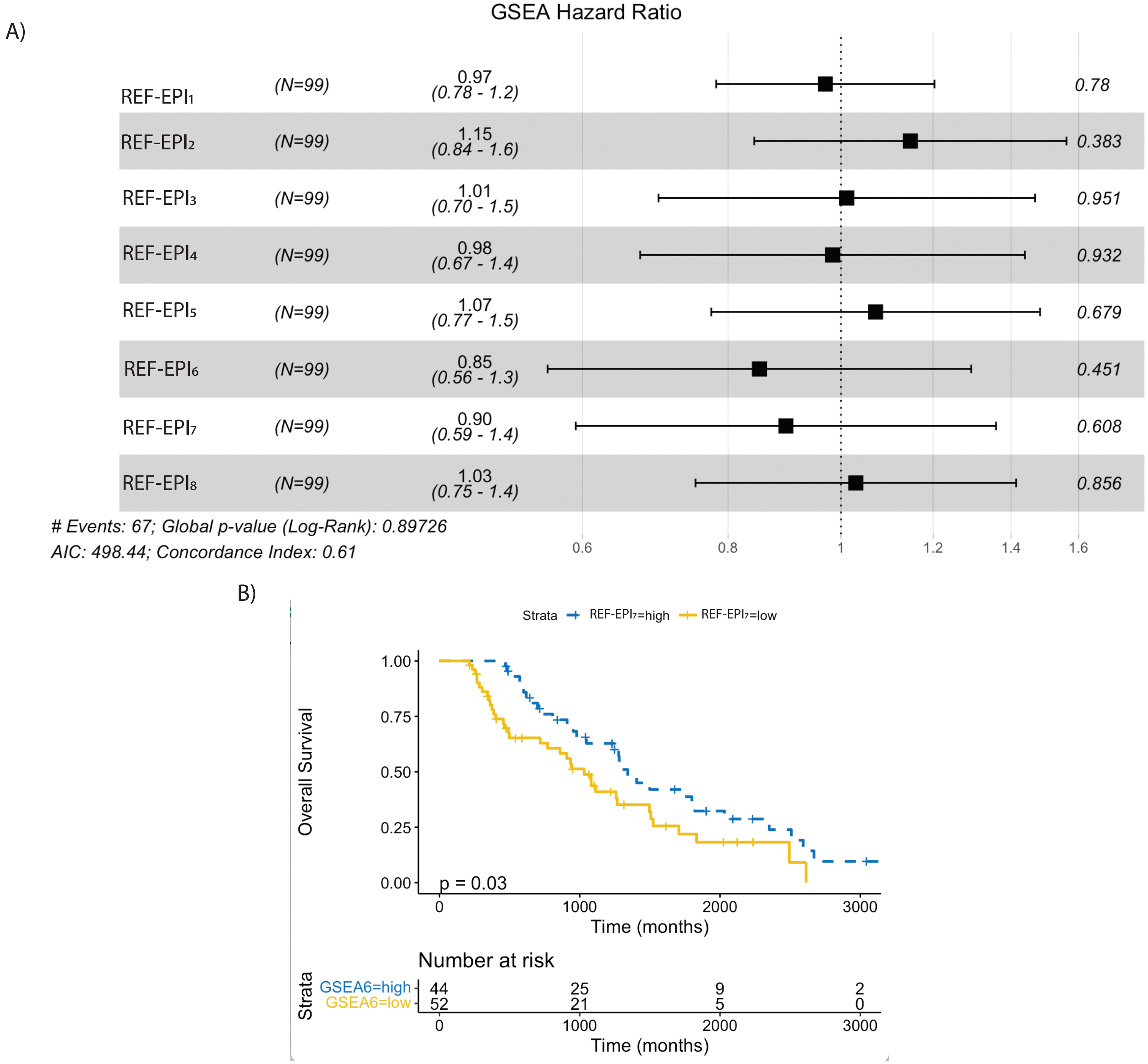
A)
GSEA Hazard Ratio
REF-EPI1 (N=99) 0.97 (0.78 - 1.2) 0.78
REF-EPI2 (N=99) 1.15 (0.84 - 1.6) 0.383
REF-EPI3 (N=99) 1.01 (0.70 - 1.5) 0.951
REF-EPI4 (N=99) 0.98 (0.67 - 1.4) 0.932
REF-EPI5 (N=99) 1.07 (0.77 - 1.5) 0.679
REF-EPI6 (N=99) 0.85 (0.56 - 1.3) 0.451
REF-EPI7 (N=99) 0.90 (0.59 - 1.4) 0.608
REF-EPI8 (N=99) 1.03 (0.75 - 1.4) 0.856
### Events: 67; Global p-value (Log-Rank): 0.89726
AIC: 498.44; Concordance Index: 0.61
0.6
0.8
1
1.2
1.4
1.6
B)
Strata
REF-EPI7=high
REF-EPI7=low
Overall Survival
1.00
0.75
0.50
0.25
0.00
p = 0.03
0
1000
2000
3000
Time (months)
Number at risk
Strata
GSEA6=high 44 25 9 2
GSEA6=low 52 21 5 0
Time (months)

Figure S4

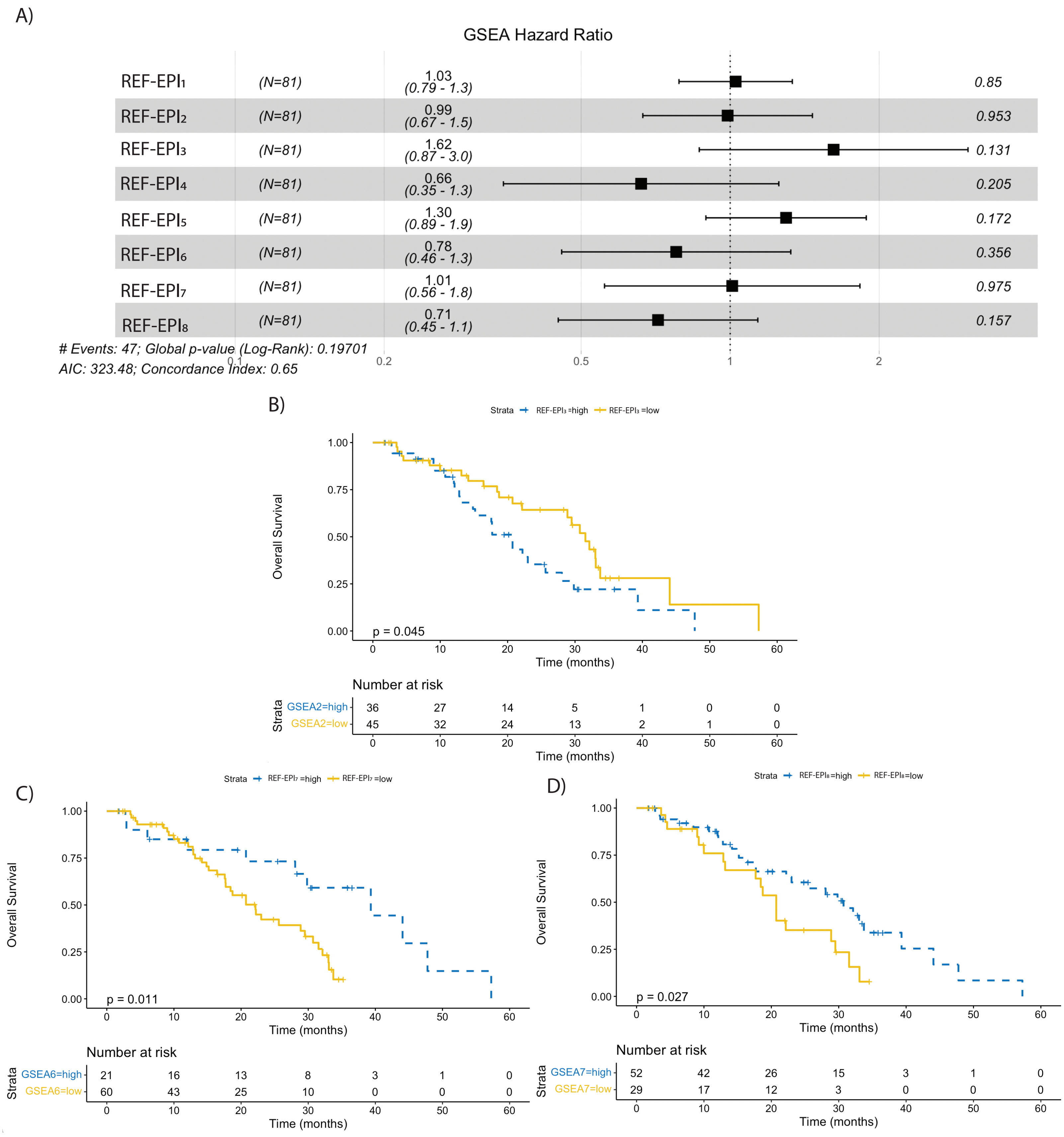
A)
GSEA Hazard Ratio
REF-EPI1 (N=81) 1.03 (0.79 - 1.3) 0.85
REF-EPI2 (N=81) 0.99 (0.67 - 1.5) 0.953
REF-EPI3 (N=81) 1.62 (0.87 - 3.0) 0.131
REF-EPI4 (N=81) 0.66 (0.35 - 1.3) 0.205
REF-EPI5 (N=81) 1.30 (0.89 - 1.9) 0.172
REF-EPI6 (N=81) 0.78 (0.46 - 1.3) 0.356
REF-EPI7 (N=81) 1.01 (0.56 - 1.8) 0.975
REF-EPI8 (N=81) 0.71 (0.45 - 1.1) 0.157
### Events: 47; Global p-value (Log-Rank): 0.19701
AIC: 323.48; Concordance Index: 0.65
0.1 0.2 0.5 1 2
B)
Strata REF-EPI3 =high REF-EPI3 =low
Overall Survival
p = 0.045
Time (months)
Number at risk
Strata
GSEA2=high 36 27 14 5 1 0 0
GSEA2=low 45 32 24 13 2 1 0
Time (months)
C)
Strata REF-EPI7 =high REF-EPI7 =low
Overall Survival
p = 0.011
Time (months)
Number at risk
Strata
GSEA6=high 21 16 13 8 3 1 0
GSEA6=low 60 43 25 10 0 0 0
Time (months)
D)
Strata REF-EPI8=high REF-EPI8=low
Overall Survival
p = 0.027
Time (months)
Number at risk
Strata
GSEA7=high 52 42 26 15 3 1 0
GSEA7=low 29 17 12 3 0 0 0
Time (months)

Figure S5

### Hallmarks of Cancer Pathway Enrichment

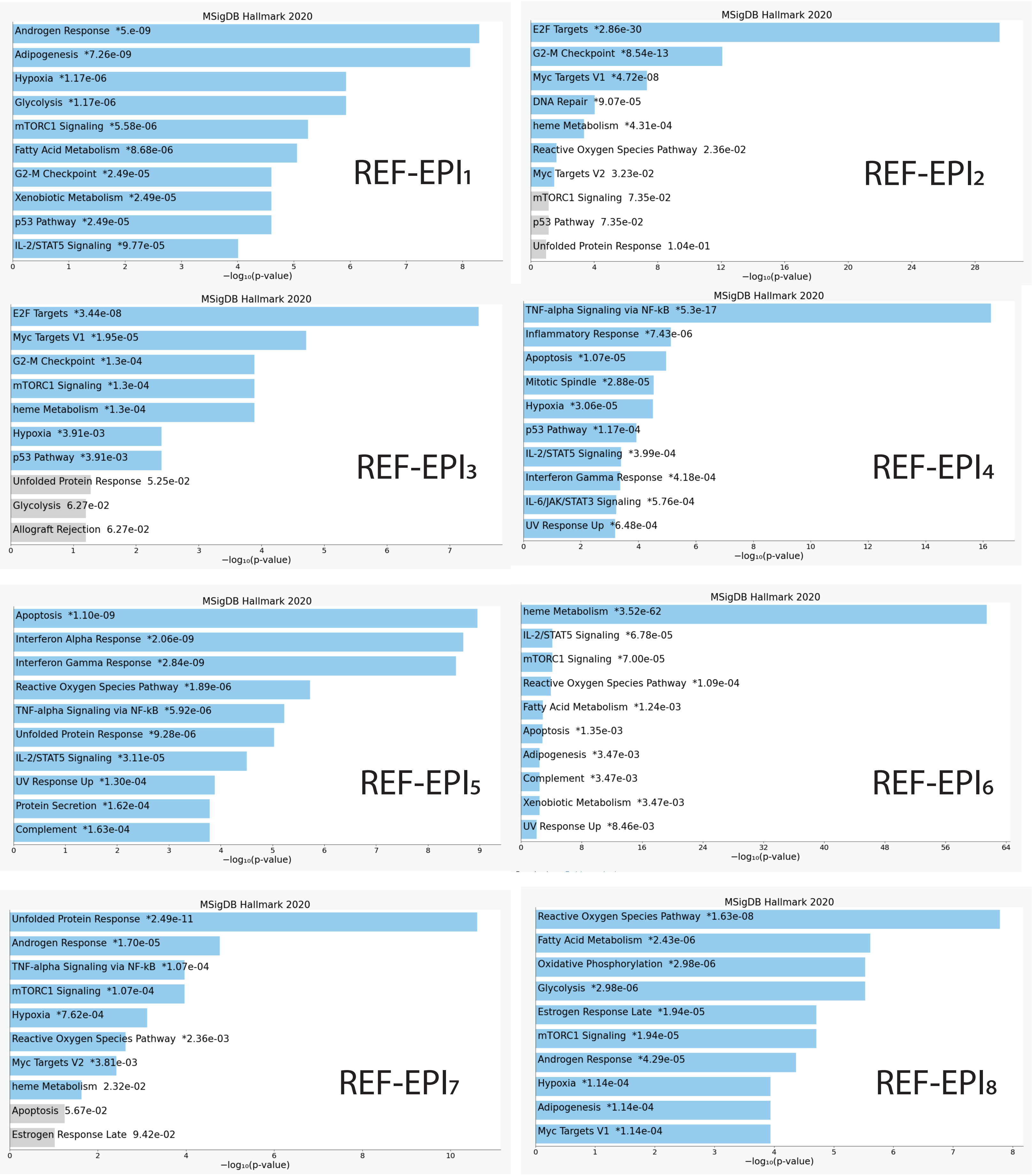

Figure S6

A)

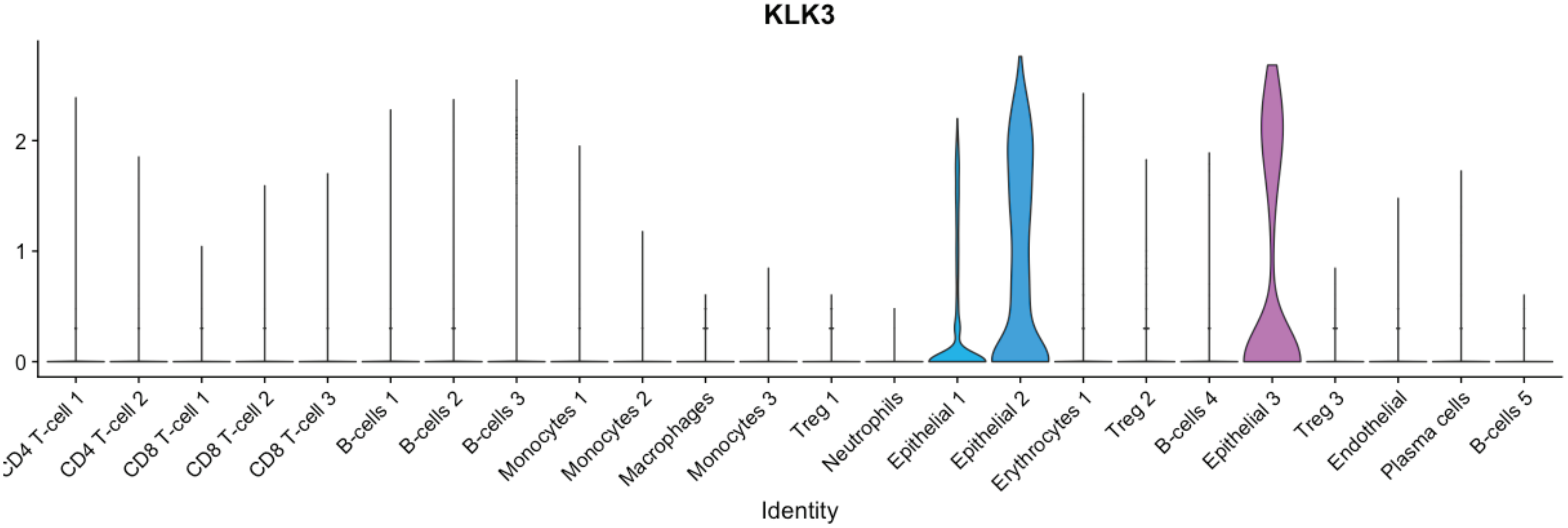
KLK3
0
1
2
CD4 T-cell 1
CD4 T-cell 2
CD8 T-cell 1
CD8 T-cell 2
CD8 T-cell 3
B-cells 1
B-cells 2
B-cells 3
Monocytes 1
Monocytes 2
Macrophages
Monocytes 3
Treg 1
Neutrophils
Epithelial 1
Epithelial 2
Erythrocytes 1
Treg 2
B-cells 4
Epithelial 3
Treg 3
Endothelial
Plasma cells
B-cells 5
Identity

B)

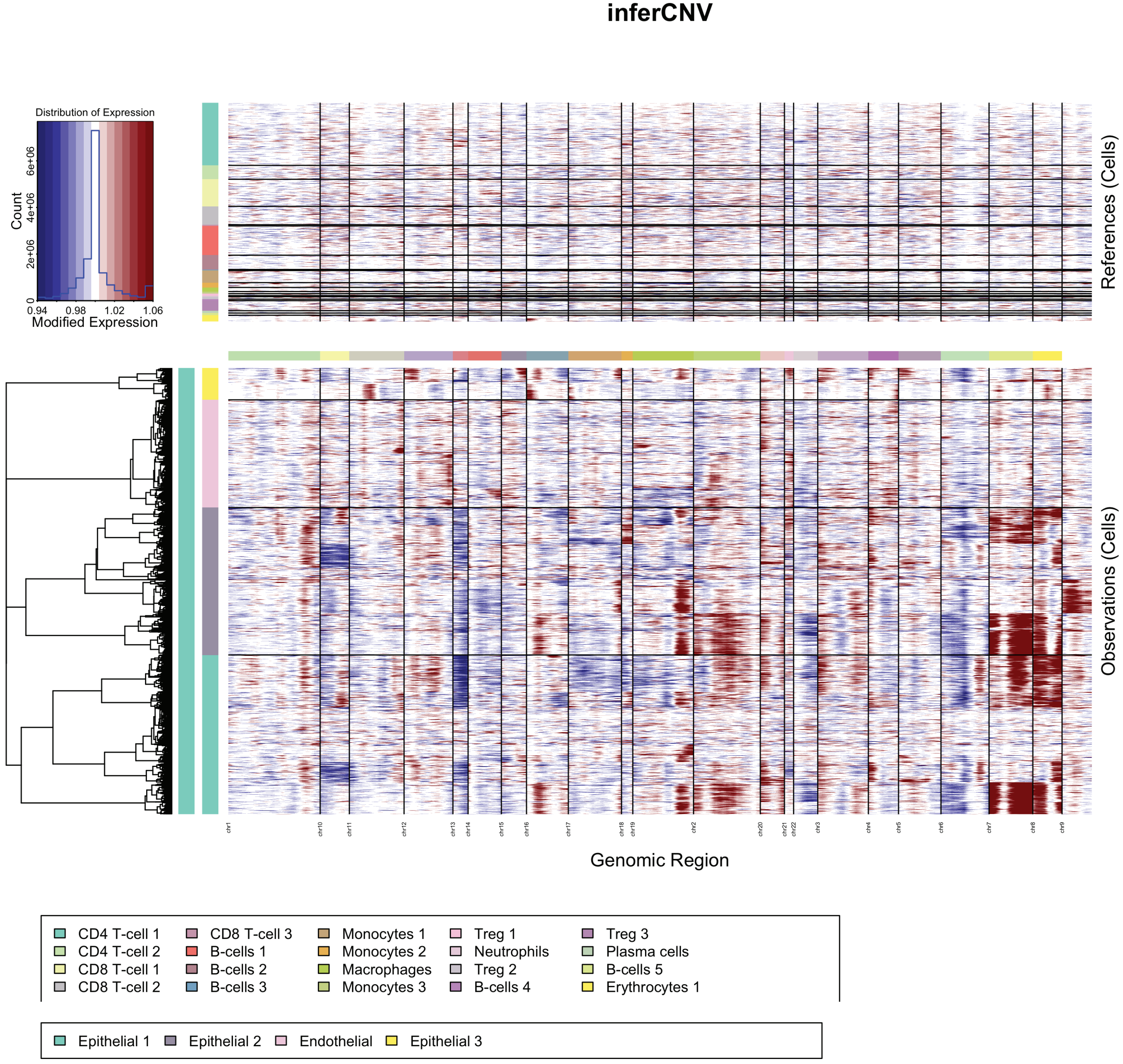
inferCNV
Distribution of Expression
Count
0
2e+06
4e+06
6e+06
0.94
0.98
1.02
1.06
Modified Expression
References (Cells)
Observations (Cells)
Genomic Region
CD4 T-cell 1
CD4 T-cell 2
CD8 T-cell 1
CD8 T-cell 2
CD8 T-cell 3
B-cells 1
B-cells 2
B-cells 3
Monocytes 1
Monocytes 2
Macrophages
Monocytes 3
Treg 1
Neutrophils
Treg 2
B-cells 4
Treg 3
Plasma cells
B-cells 5
Erythrocytes 1
Epithelial 1
Epithelial 2
Endothelial
Epithelial 3
